# Comparative single-cell analyses identify shared and divergent features of human and mouse kidney development

**DOI:** 10.1101/2023.05.16.540880

**Authors:** Sunghyun Kim, Kari Koppitch, Riana K. Parvez, Jinjin Guo, MaryAnne Achieng, Jack Schnell, Nils O. Lindström, Andrew P. McMahon

## Abstract

Mammalian kidneys maintain fluid homeostasis through the cellular activity of nephrons and the conjoined collecting system. Each epithelial network originates from distinct progenitor cell populations that reciprocally interact during development. To extend our understanding of human and mouse kidney development, we profiled chromatin organization (ATAC-seq) and gene expression (RNA-seq) in developing human and mouse kidneys. Data were analyzed at a species level and then integrated into a common, cross-species multimodal data set. Comparative analysis of cell types and developmental trajectories identified conserved and divergent features of chromatin organization and linked gene activity, revealing species- and cell-type specific regulatory programs. Identification of human-specific enhancer regions linked through GWAS studies to kidney disease highlights the potential of developmental modeling to provide clinical insight.

## INTRODUCTION

The mammalian kidney is a pivotal organ in the homeostasis of body fluids^1^. The nephron plays the predominant role in renal recovery of essential factors—proteins, metabolites, ions, and elimination of toxic compounds, while the ureteric epithelium modulates water, salt and pH balance. Loss of kidney function and end stage renal disease (ESRD) are associated with high morbidity and mortality, with patients requiring dialysis or kidney transplants, the latter being the only effective long term solution^2^. An enhanced understanding of kidney development is already facilitating the generation of disease-appropriate models for therapeutic discovery, and ultimately kidney surrogates capable of restoring kidney functions to patients with ESRD^3^.

The joint nephron and collecting system epithelial network is assembled from distinct progenitor cell types over a lengthy period of development: 10-11 day in the mouse and 26-28 weeks in the human kidney^4,5^. Interactions between nephron, ureteric, interstitial, and vascular progenitor cells (NPC, UPC, IPC, VPC, respectively) within nephrogenic niches, lead to the induction and mesenchymal-to-epithelial transition of NPCs to generate the nephrons of the kidney: approximately 14,000 in the mouse and one million in the human kidney^6,7^. Alongside the formation of nephrons, the adjacent ureteric epithelium branches and grows to generate the arborized network of the collecting system. The connection of developing nephrons to the collecting system generates a continuous renal epithelial network^8^. Single cell transcriptional studies of the mouse kidney have highlighted remarkable cellular diversity in this tubular epithelial network related to the time of nephron formation, sex, and the position of cells along the kidney’s corticomedullary axis^9–12^. Studies of the adult human kidney have revealed conservation of cell diversity and cell organization between these species^13–16^.

Much of our understanding of mammalian kidney development has centered on the mouse as the developmental and genetic model system. The concordance in developmental phenotypes in mouse models of human congenital disease support the argument that mouse and human developmental programs are deeply conserved^17–20^. Comparative developmental studies have highlighted conserved anatomies and molecular signatures in early nephron and ureteric epithelial programs^13–15,21–25^. However, while human and mouse species do share considerable similarities, they have followed separate evolutionary paths, with distinct selective pressures, for around 80 million years^26^. At a gross anatomical level, mouse and human kidneys differ in size, nephron numbers and nephron connectivity, human-specific lobulation and patterns of collecting system arborization^27,28^.

Though there has been no comprehensive comparison of mouse and human kidney developmental programs to date, direct comparisons of NPCs have identified species differences^21,29^. Single cell technologies provide new opportunities to compare molecular and cellular processes in mouse and human kidney development^30^. To this end, we constructed comparative developmental atlases detailing human and mouse transcriptomic and chromatin accessibility profiles across kidney lineages. Analysis of these data identified conserved and divergent features of chromatin organization and linked gene activity and predicted species-specific and cell-type specific regulatory features associated with kidney development and disease.

## RESULTS

### Transcriptomic and epigenomic atlases of human and mouse kidney organogenesis

To extend our understanding of conserved and divergent features of human and mouse kidney organogenesis and detail their respective cell diversity, genomic organization, and gene activity, we performed single-cell (mouse) and single-nuclear (human) RNA-seq (sc/snRNA-seq) profiling and single nuclear ATAC-seq (snATAC-seq) profiling and scrutinized gene expression and epigenetic profiles (Figure 1A). Mouse studies focused on the neonatal kidney the postnatal day of birth (p0, n=10) when the kidney is both functional and life-supporting but still undergoing substantial development: approximately 50% of the mouse complement of nephrons form after birth^31,32^. Human fetal kidneys ranged in age from 10.6-17.6 week (n=10), a period associated with the appearance of mature nephron cell types and acquisition of function; the bulk of human nephrons form in the remaining period of human kidney development to 32 weeks^22,33^ sn/scRNA-seq data were generated using 10X Chromium v2/v3, DNB-seq and Novaseq platforms. After applying quality-control metrics and eliminating potential sequencing artifacts through the Seurat V3 R package^34^ (Figure S1A, B), we obtained 52,046 nuclei with a median of 2,780 genes per nucleus for human snRNA-seq data set, and 57,431 cells with a median of 2,813 genes per cell for mouse scRNA-seq data set. To identify parallel chromatin accessibility landscapes in mouse P0 (n=4) and human 10.6-17.6 week kidneys (n=9), we performed an assay for transposase-accessible chromatin (ATAC)^35^ on single nuclei using 10X Genomics Chromium ATAC v1.1, DNB-seq and Illumina Nova-seq platforms. After quality control with the Signac R package^36^ (Figure S1C, D), we obtained 79,629 human nuclei with a median of 6,476 fragments per cell and 33,235 mouse nuclei with a median of 12,457 fragments per cell. CCA-based integration in Seurat v3 (sc/snRNA-seq) or Harmony integration (snATAC-seq) were implemented to resolve batch effects amongst different biological samples reflecting differences in 10X Chromium chemistry, sequencing platforms and the timing of sample analysis (Table S1; tabulated list of all sequencing data in this study).

**Figure 1.**
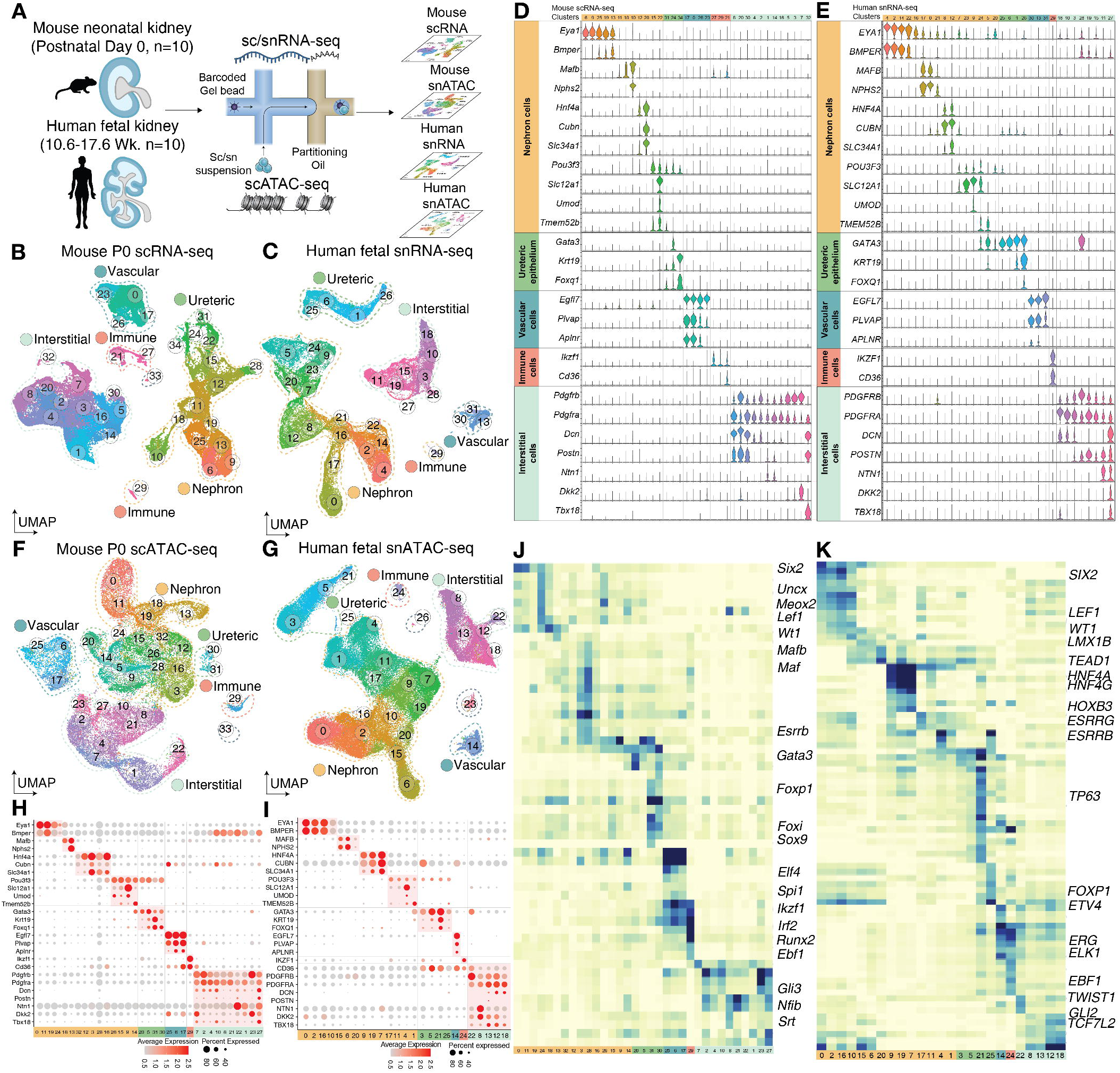
Paired single-cell/nuclei transcriptomic and open chromatin accessibility atlases of the developing human and mouse kidney. A, Overview of methodology, tissue source and single-cell/nuclei sequencing. B-C, UMAP projections of developing mouse kidney scRNA-seq (B) or human kidney snRNA-seq dataset (C) annotated to five major cell-type groupings (dashed lines): nephrogenic – yellow, ureteric – green, vascular – blue, immune – red, interstitial – light green. D-E, Violin plots showing harmonized gene expression markers of mouse (D) or human (E) sc/snRNA-seq data arranged by group and cell type, cluster numbers at top. F-G, UMAP projections of developing kidney snATAC-seq data, mouse (F) or human (G) annotated as in (B-C). H-I, Dot plots of gene activity in developing mouse (H) or human (I) kidney snATAC-seq. J-K, Heat maps of TF binding motif prediction in developing mouse (J) or human (K) kidney snATAC-seq. *See also Figure S1*

Unsupervised UMAP (Uniform Manifold Approximation and Projection) clustering of the four datasets in both data modalities highlighted cells of the five major kidney lineage compartments: nephrogenic (N), ureteric epithelium (UE), vascular (V), interstitial (INT), and immune (IMU) (Figure 1B, C; sc/snRNA-seq, 1F, G; snATAC-seq). For sc/snRNA-seq data, cell-type clusters were stratified based on: (1) cell-type marker genes of well-accepted gene expression signatures; (2) the high cell-type enrichment of gene expression for individual clusters; and (3) harmonizing concordance in paired human and mouse clusters illustrated in the violin plots (Figure 1D, E). To identify clusters in the snATAC-seq data, gene expression levels were inferred by generating a “gene activity score”, summing snATAC-seq fragments per cell within the gene body together with upstream (2kb) promoter regions^36^. Comparable cell-type identities were predicted in the developing mouse and human kidney based on gene activity scores (Figure 1H, I).

To identify developmental regulators, we examined the enrichment of transcription factor (TF) binding motifs within each nucleus-grouping using chromVAR^37^ and JASPAR data^38^. The predicted binding site enrichments showed a concordance with cell-type specific regulators of gene expression that are known to drive renal developmental programs (Figure 1J, K). Shared TF expression specific to distinct lineages and cell-types within the mouse and human sc/snRNA-seq data correlated well with predictions for TF engagement from analysis of enriched binding motifs within the snATAC-seq datasets.

In addition to canonical renal cell identities, mouse scRNA-seq cluster 33 and snATAC-seq cluster 33 likely represents Schwann cell precursors of the migratory neural crest cell lineage expected to associate with the renal sympathetic neuron innervation and distinguished by enriched expression of key TFs *Sox10*^39^, and *Foxd3*^40^ and other genes associated with Schwann cell types e.g., *Plp1*, *Mpz*, and *S100b*^41^ (Figure S1E). Further, the human snATAC-seq cluster 26 is tentatively identified as autonomic neuron precursors of a neural crest origin based on differential gene activity of *SRRM4*^42,43^, *PHOX2B*^44,45^, and *HANDS2*^46,47^ (Figure S1F).

### Progenitors and functionally maturing cells in nephron and ureteric epithelium compartments

To determine the range of nephron and ureteric cells captured from each species for their respective developmental times, the nephrogenic and ureteric lineages were subset and progenitors identified as *EYA1^+^/Eya1^+^* NPCs and *RET^+^/Ret^+^*UPCs (Figure 2B, C). Human and mouse transcriptional profiles correlated closely (Figure S2A) and shared cell-type specific TF activity across developmental landscapes (Figure S2B).

**Figure 2.**
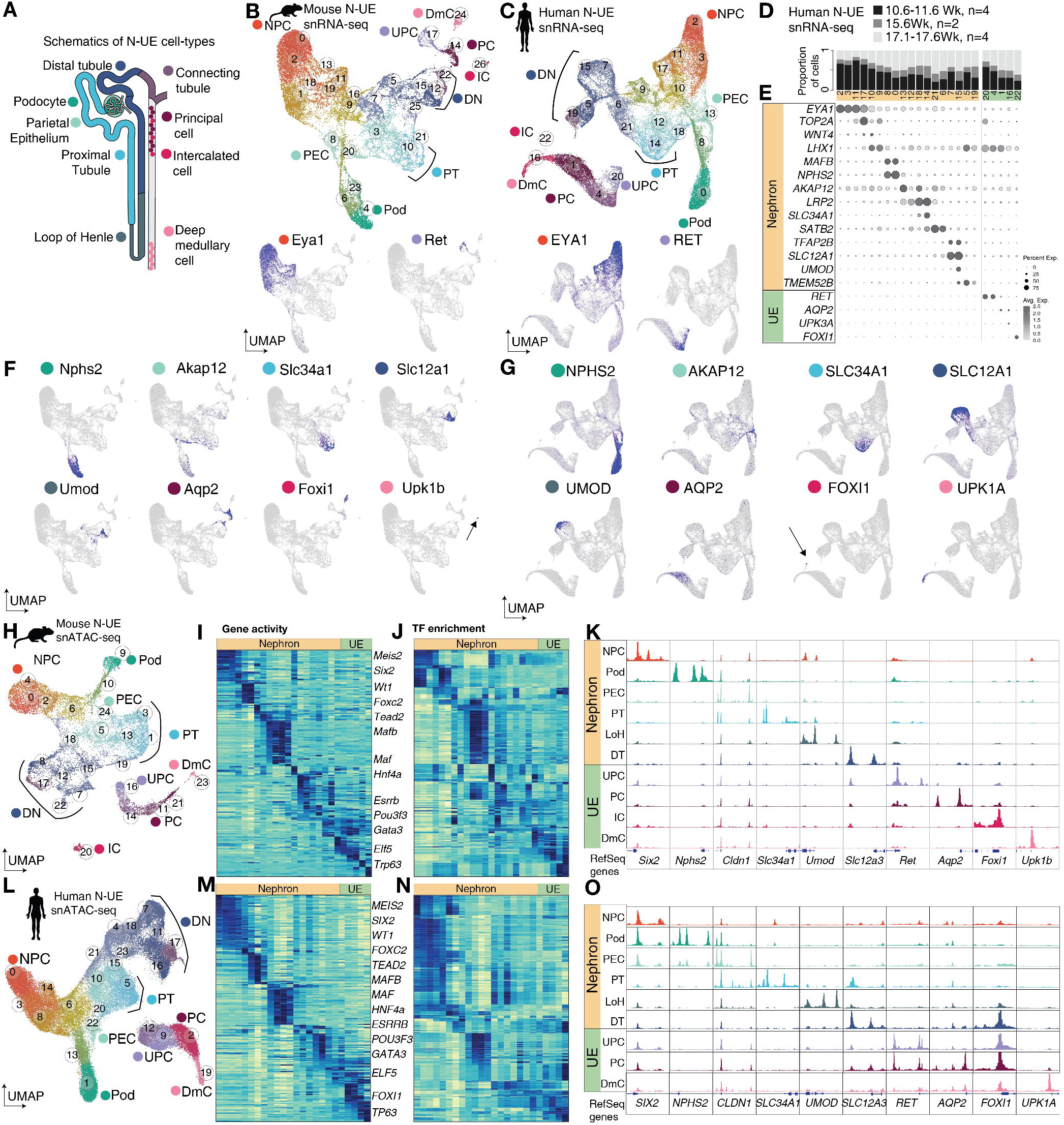
Progenitor and differentiated the nephron (N) and ureteric epithelium (UE) cell-types in the developing human and mouse kidney. A Schematic illustration and color-coding of regional diversity for N and UE cell types. Color code reflects diversity in UMAP clusters and violin plots below. B-C, UMAP projections showing sub-clustering analysis of developing mouse N-UE scRNA-seq (B) and human N-UE snRNA-seq (C). D-E, Stacked bar plot of per cent of cells with differential contribution in the human N-UE snRNA-seq clusters (D) and dot plot representation of gene expression markers (E) F-G, UMAP projection of signature marker gene expression for terminally differentiated cell-types in the mouse scRNA-seq (F) and the human snRNA-seq (G). H-L, UMAP projections showing subclustering analysis of developing mouse N-UE snATAC-seq (H) and human N-UE snATAC-seq (L). I-M, Heat maps of snATAC-seq derived-gene activity score of mouse (I) and human (M) N and UE cell-types. J-N, Heat maps of TF binding motif recovery of mouse (J) and human (N) N and UE cell-types. K-O, Genome browser views of aggregated pseudo-bulk ATAC-seq displaying cis-regulatory regions including promoters and upstream elements of mouse (K) and human (O) N-UE cell-type marker genes. *See also Figure S2*

To examine terminal cell-type representation in the developing nephron and ureteric compartments, we correlated signatures at P0 to those in the spatially resolved adult mouse kidney^10^ (Figure 2 A, F, G). Clear relationships were evident in the Pearson correlation heatmaps though markedly less cellular diversity was observed within the loop of Henle in the p0 mouse kidney. Marker genes for tDL and SDL, loop of Henle cell-types (e.g., *Slc14a2, Gdf15, Fst, Pitx2, Bcl6, and Sptssb*) were co-expressed in few cells in a minor cluster (cluster 25) (Figure S2C). In the human kidney, expression of key signature marker genes expressed within mature human cell types^105^ distinguished major N and UE cell-types in terminally located clusters in individual UMAP plots (Figure 2H, I).

In line with expectations, we observed a proportionally higher representation of progenitor populations within early 10.6-11.6 week human kidney samples (Figure 2D, E; *EYA1^+^* NPCs in cluster 11 and 17; *RET^+^* UPCs in cluster 20) and more differentiated cells in older 17.1-17.6 week kidneys (*NPHS2^+^* podocytes in cluster 0), *LRP2^+^* proximal tubule cell types (clusters 12, 13, 14, 18, 21), *UMOD^+^* loop of Henle cells (cluster 15) and *FOXI1^+^*intercalated cells (cluster 22). Whole kidney mRNA-seq studies over a similar time course have highlighted corresponding trends^22^.

A comparative analysis of nephron and ureteric cell-types using snATAC-seq datasets (Figure 2K, O) identified comparable cell types predicted by gene activity score and TF motif enrichment analysis (Figure 2L-M, 2P-Q). Differentially accessible open chromatin regions for marker genes showed agreement in accessibility of promotor regions for differentiated cell types and of open chromatin flanking 5’ and intronic putative enhancer elements (Figure 2N and R).

In summary, N-UE sub-clustering demonstrated expected features of a dataset comprising transcriptomic and chromatin profiling incorporating a transition from progenitor to terminally differentiated mouse and human kidney cell-types. Tabulated lists of the differentially expressed genes (DEGs), differentially accessible regions (DARs), and differential TF binding predictions for each cluster can be viewed in Table S2.

### Predicted developmental trajectories demonstrate broad similarities and distinct human-specific features

To directly compare mouse and human differentiation trajectories for the nephrogenic and ureteric programs, we employed RNA velocity^48^ to predict the future states of N and UE lineage cells through the time derivative of gene expression states (Figure S3A). In the UE lineage, velocity analysis highlights a flow from UPC to differentiated principal cells (PC) cells. In the human and mouse N lineage, committed *WNT4/Wnt4^high^* cells show an expected flow in the direction of more differentiated cell types, as well as trajectory towards an *EYA1/Eya1^high^*NPC population. Dynamic studies of mouse *Wnt4+* cell types^49^ and single-cell studies of the developing mouse kidney^50^ have demonstrated occasional plasticity in this commitment step. The findings here suggest a high level of developmental flexibility in response to inductive signaling at the outset of human nephrogenesis.

To generate comprehensive N lineage compartments, cells from the *PAX8^+^/Pax8^+^*NPCs to terminally differentiated N-lineage cells were selected for re-clustering, to exclude potential artifacts caused by the proliferative cells in RNA velocity analysis. To this end, the three lobular shapes in the re-clustered UMAPs clearly indicated lineage-specific TF gene activity signatures —*MAFB/Mafb*, *HNF4A/Hnf4a*, and *POU3F3/Pou3f3*, further assigned into podocyte-, PT-, and distal nephron (DN)-lineages, respectively (Figure 3A; Table S3 for differentially expressed genes). In addition, we performed in silico lineage separation by Hox gene profiling. N and UE lineages arise at different times and positions in the mammalian embryo^51^. Temporal and spatial differences are evident in differential expression of Hox genes, which distinguishes amongst distal nephron (*Hoxd10/11* positive) and UE-derived (*Hoxd10/11* negative) cell types with very similar transcriptional programs and cellular properties^10^. Isolating and re-integrating *HOXD10/11* or *Hoxd10/11*-positive cells and nuclei which show UE features allowed the assembly of complete N lineage objects for each species (Figure S3B).

**Figure 3.**
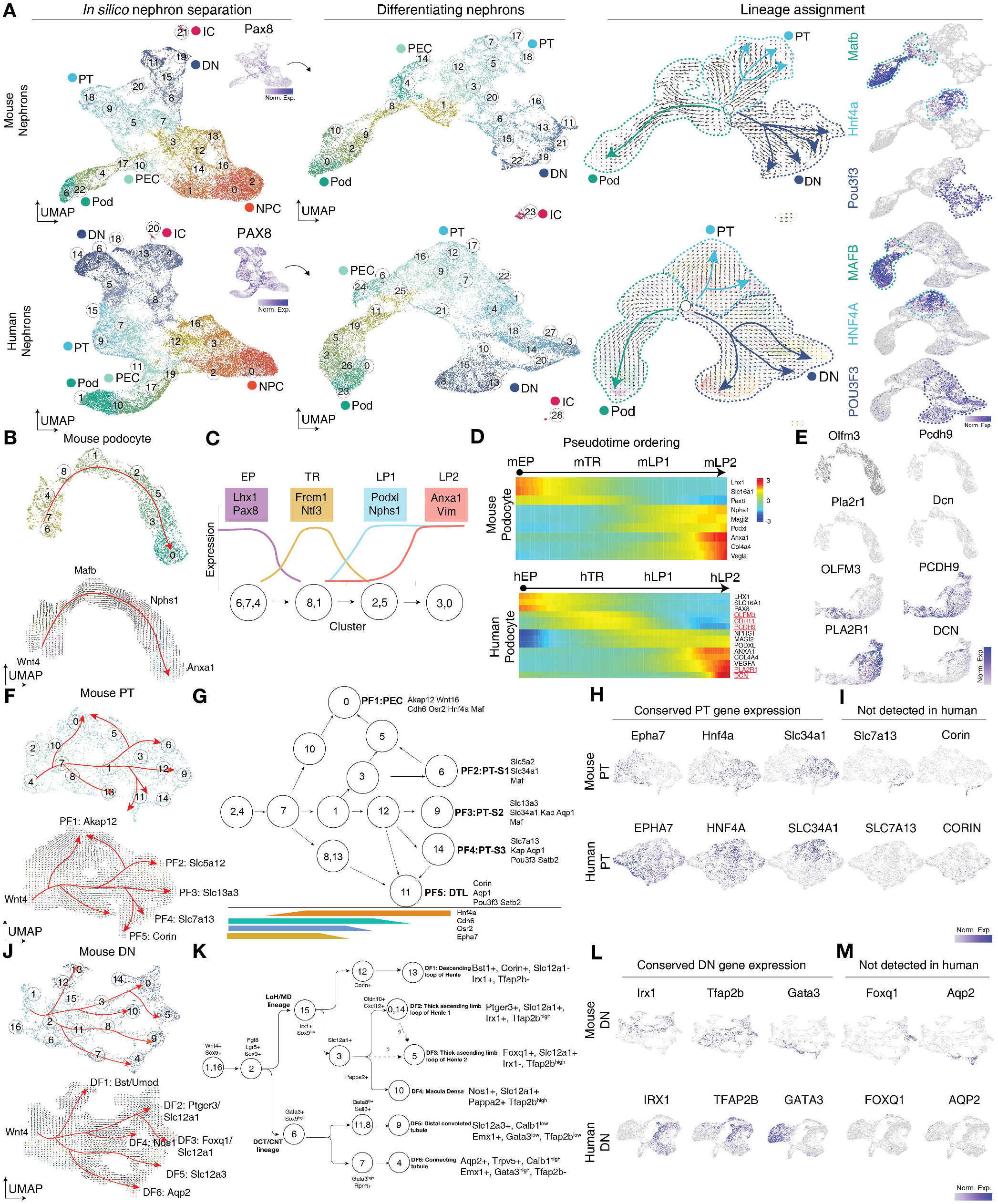
Prediction of the human and mouse nephron trajectories showed broad similarity but distinct human-specific features in development and differentiation outcome. A, Subclustering analysis and lineage assignment strategy of *PAX8^+^/Pax8^+^*differentiating nephrons and NPC derivatives. B, UMAP projection of mouse podocyte (top) and RNA velocity analysis (bottom) with cell-type annotation. C, Schematic diagram of mouse podocyte lineage specification. D, Pseudotime ordered mouse (top) and human (bottom) podocyte dynamic gene expression program, human-specific gene expression marked red. E. human-specific gene expression *OLFM3*, *PCDH9*, *PLA2R1* and *DCN*, and no mouse orthologous gene expression. F, UMAP projection and RNA velocity analysis on mouse PT lineage cells. G, Schematic diagram of mouse PT lineage specification. H, Conserved gene expression in mouse and human PT cells, *EPHA7*, *HNF4A*, and *SLC34A1*. I, Gene activity of mouse PT-S3 and DTL marker, *Slc7a13* and *Corin*, and no human ortholog expression. J, UMAP projection and RNA velocity analysis on mouse DN lineage cells. K, Schematic diagram of mouse DN lineage specification. L, Conserved gene expression in mouse and human DN cells, *IRX1*, *TFAP2B*, *EMX1*, and *GATA3*. M, Gene activity of mouse TAL2 and CNT marker, *Foxq1* and *Aqp2*, no human ortholog expression. *See also Figure S3* NPC, nephron progenitor cells; Pod, podocyte; PT, proximal tubule, PEC, parietal epithelial cell; DN, distal nephron; IC, intercalated cell; EP, early podocyte; TR, transient expression genes; LP, late podocyte; DTL, descending thin limb of LoH; DCT, distal convoluted tubule; CNT, connecting tubule.

### Juxtaposition of the human and mouse podocyte development and differentiation outcomes

To predict gene expression changes in conjunction with human and mouse nephrogenesis, we performed pseudotime ordering of the RNA velocity-inferred lineages in Monocle2^53^. Mouse podocyte trajectories were classified into four groups (Figure 3B, C): 1) early podocyte-restricted activity (EP; *Lhx1*, *Pax8*, and *Slc16a1*), 2) transient expression genes (TR; *Frem1*, *Ntf3*), 3) activated early and remaining high through the trajectory (LP1; *Nphs1*, *Magi2*, and *Podxl*), and 4) late podocyte gene expression upregulated at the terminal state (LP2; *Anxa1*, *Col4a4*, and *Vegfa*). Mouse and human podocyte programs showed largely conserved dynamic gene expression programs and a good agreement with earlier observations of the temporal progression of human podocyte development^55^ (Figure 3D). However, several human TR genes (e.g., *OLFM3* and *PCDH9*) and some terminal markers of human podocyte differentiation (e.g., *PLA2R1* and *DCN*) showed little or no equivalent expression in the mouse podocyte lineage (Figure 3E). Further, analysis of snATAC-seq profiles for the human specific TR genes *OLFM3*, *PCDH9*, *PLA2R1*, and *DCN* indicated open chromatin profiles for the gene body and potential upstream regulatory regions for the human genes that were absent from the loci of their mouse orthologs (Figure S3C). Thus, species specific differences in gene regulations are likely to account for differences in gene expression in human and mouse podocyte developmental programs.

### Dynamic gene expression programs in the mouse and human PT and DN

Next, we extended mouse and human comparisons of lineage progression to PT (*HNF4A^+^/Hnf4a^+^*) and DN (*POU3F3/Pou3f3^+^)* sub-compartments, curating gene expression signatures first on mouse datasets, then cross-referencing conservation of orthologous genes in human developmental trajectories. Along the *Hnf4a^+^* PT trajectory, robust *Hnf4a* upregulation follows a transient activation of a number of genes including *Lhx1*: *Epha7*, *Osr2* and *Cdh6* (Figure 3G). Distinct expression of genes encoding solute carrier proteins (“Slc” genes) specific to individual PT segments could be identified in distinct terminal UMAP clusters, e.g., *Slc5a2* (S1; cluster 6; proximalizing fate (PF) 2), *Slc13a3* (S2; cluster 9; PF3), *Slc7a13* (S3; cluster 14; PF4). Proximally, PTs connect to *Akap12*/*Wnt16* parietal epithelial cells (PECs, cluster 0; PF1) and distally to *Corin*-expressing cells of the short descending thin limb of the loop of Henle (SDL; cluster 11; PF5) (Figure 3F). The data distinguish a proximal-distal axis of key regulatory factors (*Hnf4a*, *Irx1* and *Pou3f3*) directing regional patterning of the S-shaped body, a key intermediate in nephron construction^23^ (Figure S3D). Human PT trajectories were largely comparable to the mouse, as the human PT cells showed the conserved gene expression key PT lineage-specific factors, *CDH6*, *EPHA7* and *HNF4A* (Figure 3H), and PT-S1/S2 markers, e.g., *SLC34A1*, (Figure 3I). Human data has highlighted *HNF4A*-positive subpopulation of human PECs^52^.

Analyzing mouse *Pou3f3*^+^ cells revealed a transient expression of the distal-patterning associated genes *Fgf8* and *Lgr5*^54,56^ preceding the emergence of a six terminally differentiated DN cell-types based on unique gene expression signatures (Figure 3H, J): 1) *Bst*^+^/*Umod*^+^ descending thin limb of LoH (DTL; cluster 13; distalizing fate (DF) 1), 2) *Ptger3*^+^/*Slc12a1*^+^/*Umod*^+^ thick ascending limb of LoH type 1 (TAL1; cluster 0; DF2), 3) *Foxq1*^+^/*Slc12a1*^+^/*Umod*^−^ TAL type 2 (TAL2; cluster 5; DF3), 4) *Slc12a3*^+^/*Calb1*^+^ distal convoluted tubule (DCT; cluster 9; DF5), 5) *Aqp2*^+^/*Calb1*^+^ connecting tubule (CNT; cluster 4; DF6), and 6) *Pappa2*^+^/*Nos1*^+^ macula densa cells (MD; cluster 10; DF4). Additionally, the *Bst*^+^ DTL cells (cluster 13) includes two distinct sub-populations with *Aqp1*/*Pitx2* and *Slc6a12*/*Bcl6* gene activity (Figure S3F), suggestive of juxta-medullary thin descending limb of LoH (tDL) and cortical short descending limb of LoH (SDL), respectively.

Next, we investigated each individual DN cell-type program (Figure 3K). Distalizing nephrons in the S-shaped body stage were largely demarcated by *Irx1*^+^/*Sim2*^+^/*Sox9^low^*LoH/MD precursors and *Sox9^high^* DCT/CNT precursor cell-types. The Bst1/*Umod*^+^ DTL clusters showed the specific enrichment of *Satb2*, *Fmod*, *Corin*, and *Wnt7b* gene activity along the trajectory (Figure S3F). Of note, *Corin*^+^ expression was detected in PT-S3 and DTL, implying the coordinated contribution of DN and PT cells to LoH formation. *Ptger3*^+^/*Slc12a1*^+^ TAL1 and *Nos1*^+^/*Pappa2*^+^ MD cells showed common expression of *Cldn10* and *Cxcl12* (Figure S3G), Examining the DCT/CNT trajectories identified *Sox9* gene activity in the early DCT/CNT program, *Sall3* expression along the DCT trajectory and the sequential expression of *Bmp2*, *Gata3*, *Rprm*, *Aldh1a3*, *Crybb1* along the CNT trajectory (Figure S3H, I). *Emx1* expression was common to both late DCT/CNT programs. The RNA velocity analysis implies *Foxq1*/*Slc12a1*^+^ TAL2, a recently identified TAL subpopulation^12^, shares a developmental origin from precursors generating TAL1 and MD.

Comparing mouse with human DN trajectories, distinct combinatorial patterns of expression of the TFs *IRX1/Irx1*, *TFAP2B/Tfap2b*, *EMX1/Emx1*, and *GATA3/Gata3* were shared between human and mouse terminal DN cell-types (Figure 3L). Most representative DN-subsegment markers also showed comparable conserved expression patterns (e.g., *SLC12A1*, *PTGER3*, *SALL3*, *SLC12A3*, *BMP2*, *GATA3*, *PAPPA2*, and *NOS1*). However, human data exhibited reduced cellular diversity likely reflecting an immature state of human distal nephron development up to 17 weeks of human kidney development (Figure 3M).

### Conserved and divergent molecular features through integration of human and mouse single cell datasets

To analyze more directly conserved and divergent cell-type specific gene expression and chromatin accessibility features, we integrated equivalent human and mouse cell-types from sc/snRNA-seq and snATAC-seq of the N and UE lineage into a single, unified cross-species pseudo-multimodal dataset. A step-wise integration was performed using a Seurat V3 CCA-based approach^57^ (See Methods) with a total of 129,643 N-UE single cells/nuclei and data superimposed in an inclusive landscape of 25 integrated clusters (Figure 4A). Each integrated cluster harmoniously comprised both mouse and human profiles despite species differences and difference in data modality (snRNA-seq versus scRNA-seq), except for minor variability within snATAC-seq clusters 12,15,17 and 24) (Figure S4A).

**Figure 4.**
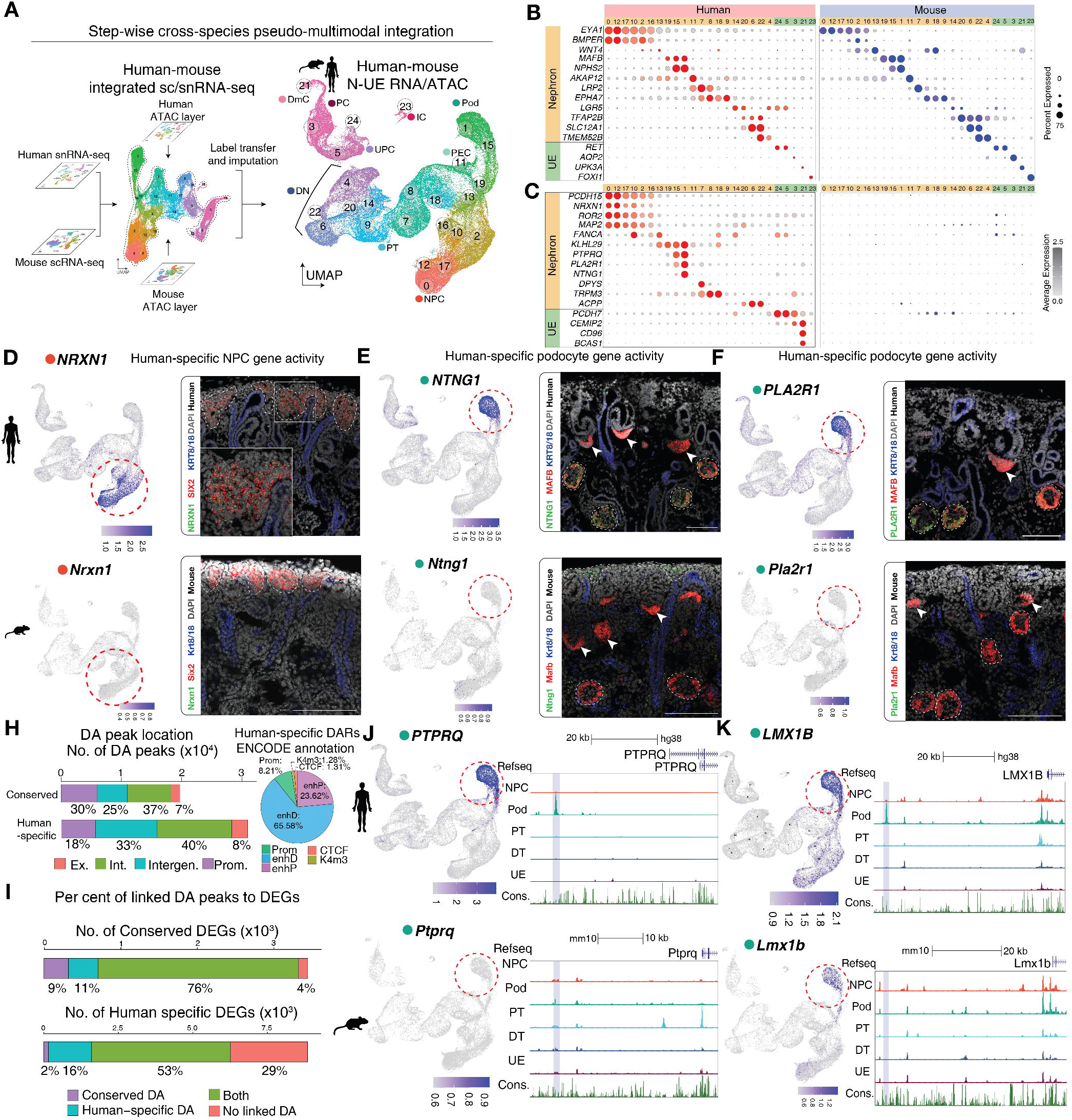
Conserved and human-enriched molecular features within the nephron and ureteric epithelium of the developing human and mouse kidney. A, Schematic and UMAP project of cross-species pseudo-multimodal integration of developing human and mouse developing kidney nephron and ureteric epithelium transcriptome and open chromatin accessibility. B-C, Cross-species dot plots of commonly expressed (B), and human-enriched (C) gene expression markers within corresponding human/mouse cell-type clusters. D-F, UMAP projection in integrated sn/scRNA-seq data and RNAScope in situ hybridization validations in human 17.6Week and mouse P0 kidney of human-only expressed genes *NRXN1*/*Nrxn1* (D), *NTNG1*/*Ntng1* (E), *PLA2R1*/*Pla2r1* (F), For E,F – arrowhead: S-shaped body, white dashed lines: glomeruli. G, Bar plot of per cent of differentially accessible regions (DARs) location in conserved and human-specific DARs. H, Bar plot of per cent of conserved and human-enriched differentially expressed genes (DEGs) associated with conserved DARs and/or human-specific DARs. I-J, UMAP and genome browser views of human-enriched DEG-human-specific DAR pair example of *PTPRQ* (I) conserved-DEG-human-specific DAR pair example of *LMX1B* (J). *See also Figure S4 and S5*

Human and mouse cell-type co-clusters showed specificity in the expression of well-accepted cell-type marker genes examining scRNAseq and snRNA-seq, though expression levels varied depending on the profiling modality (Figure 4B), and in “gene activity” predicted from snATAC-seq data (Figure S4B). Table S5 tabulates cell-type specific gene expression, computed gene activity scores, and open chromatin regions for each human/mouse integrated cluster. By intersecting human and mouse cell-type DEGs within a cluster, we catalogued 1,390 conserved, differential gene expression signatures between mouse and human (p.adj < 0.05). Using a stringent criterion for specificity (avg.FC >0.5, pct.diff > 0.3, p.adj < 0.05), this group was reduced to 352 conserved cell-type and species specific DEGs (Table S6). Furthermore, by excluding genes complementary to mouse and human DEGs, we generated a “human-enriched” cell-type DEG set of the 885 genes (avg.FC >0.5, pct.diff > 0.3, p.adj < 0.05) (Table S6).

This gene cohort comprised 229 human genes without mouse orthologs (e.g., zinc finger proteins : *ZNF730*, *ZNF385D*) (Figure S4C), a strong validation of the approach (Fisher’s exact test: *p-value* 1.526e-09), and 656 human genes with no evident expression of their matching mouse orthologs in a specific cluster (Table S4), e.g., the human NPC-specific *PCDH15* gene activity, previously identified in previous bulk mRNA-seq study^24^. Secondary validations using in situ hybridization confirmed human-specific mRNA of *NRXN1*, *ZMAT4* (NPC), *NTNG1*, and *PLA2R1* (podocytes in mature glomeruli) (Figure 4D-F, *ZMAT4*; Fig S4D). Notably, forty four TFs with known-mouse orthologs (e.g., *NR3C2*, *NPAS2*; Figure S4E) were identified in this comparative analysis.

Importantly, patterns of gene expression can differ between species even though global expression profiles may appear similar analyzing at the whole organ level^58^, a confounding factor in bulk-RNA-seq and microarray studies. As an example, *UNC5B* was expressed in human NPCs whereas its mouse counterpart *Unc5b* was absent from NPCs but expressed in multiple differentiating clusters including podocytes^24^ (Figure S4F). Conversely, in the nephron lineage, *Eya4* and Fgf1 were expressed in mouse-NPCs but expression of their human orthologs was confined to podocytes (Figure S4F). A total of 164 genes showed human-enriched gene expression differing from their mouse counterparts across clusters (Table S6). (e.g., *Spink1* – mouse PT/*SPINK1* – human DmC; *Pde4b* – mouse podocyte/*PDE4B* – human UE cells) (Figure S4F).

To predict potential functional outcomes to genes with human-enriched divergent gene expression, gene ontology (GO) analysis was performed via EnrichR^59,60^. Several biological process (BP) terms were significantly over-represented in the human data including “nervous system development (GO:0007399) (e.g., *NRXN1*, *NPAS2*, *NRG1*) and “cell-cell adhesion via plasma-membrane adhesion molecules” (GO:0098742) (e.g., *LRRC4C*, *CDH7*, *CDH9*) (Figure S4G). Interestingly, a fraction of these genes (273 of 656, 41.62%) have been classified under the “membrane part” category in Jensen COMPARTMENTS database (Binder et al., 2014; Fisher’s exact test adjusted p-value 1.01e-05, considering 6,384 “membrane part” genes amongst 18,329 genes in the Jenson COMPARTMENT library), suggesting the potential for novel intercellular interactions in human kidney development.

To examine conserved and divergent open chromatin accessibility, differentially accessible regions (DARs) were determined for each integrated snATAC-seq cluster, and the human or mouse genomic coordinates transferred into the other species reference genome using LiftOver^62^. Among the 50,592 human cell-type specific DARs (avg.FC > 0.1, p.adj < 0.05), 19,702 (38.94%) showed “conserved” open chromatin with mouse cell-type specific DARs mapping at equivalent positions in the mouse reference genome (Table S6). The majority, 30,890 (61.08%), were identified as “human-specific” DARs showing no orthologous mouse cell-type specific DARs in the corresponding syntenic regions to the human cell-type specific DARs (Figure 4H). Human-specific peaks were more likely located within distal intergenic or intronic regions compared to common peaks consistent with variability associated with cis regulatory elements (cCREs), where these regions mostly associate with ENCODE-listed enhancer-like elements (Figure 4H). In predicting human-only cis regulatory elements (cCREs), we focused on human-specific DARs listed in the ENCODE registry^63^, a total of 19,808 human-specific open chromatin regions in our dataset (Table S6).

To harness open chromatin regions to relevant regulatory domains of neighboring genes, we annotated the conserved and human-specific DARs using GREAT ^64^. Interestingly, human-specific open chromatin regions were associated with both conserved and human-enriched gene activity (Figure 4I). More than half (63.50%, 562/885 genes) of the human-specific DEGs had at least one human-unique genomic region, implicating these regions in human specific cis-regulatory functions^65,66^. For example, *PTPRQ* (Figure 4J) and *DLG2* (Figure S5A) showed human-specific open chromatin regions and human-specific gene expression in podocytes, consistent with earlier reports^67,68^.

Enhancer redundancy during evolution has been associated with an increased precision in controlling gene expression and phenotypic robustness^69,70^. We examined divergent human-specific DARs around genes with conserved gene expression in mouse and human co-clustering to identify potential examples of enhancer evolution dissociated from de novo cell type specific expression. Interestingly, 77.52% (273 of 352 genes) of human-mouse cell-type specific genes with similar expression amongst cell types were associated with at least one human-specific cell-type DAR. For example, *LMX1B*/*Lmx1b*, encoding a key transcriptional regulator of podocytes, and *ZBTB7C*/*Zbtb7c*, and *CLIC5*/*Clic5* all showed human-specific open chromatin within intronic regions not represented in the mouse snATAC-seq podocyte data (Figure 4K; Figure S5B).

### Cataloguing human putative CREs by prioritizing snATAC-seq-derived open chromatin regions and incorporating a human kidney disease context

To improve our interpretation of open chromatin regions in snATAC-seq data, we prioritized putative cis-regulatory elements (CREs) from identified cell-type DARs in a two-step computational workflow: 1) distinguishing putative CREs by leveraging integrated data modality in our pseudo-multimodal datasets; 2) cross-referencing human GWAS SNP loci relevant to human kidney diseases (Figure 5A).

**Figure 5.**
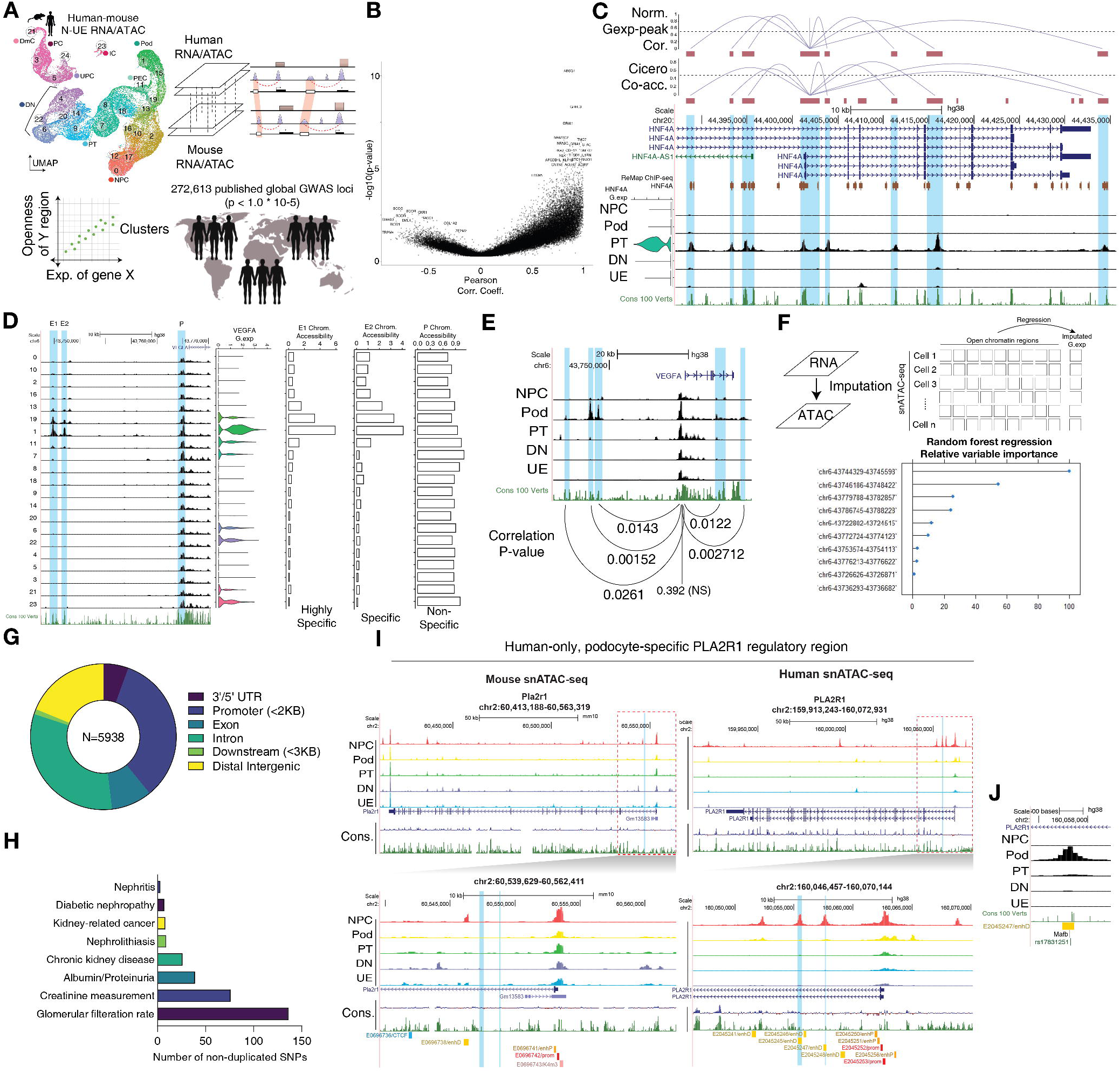
Cataloguing human putative CREs by prioritizing snATAC-seq-derived open chromatin regions and incorporating human kidney disease context. A, Schematic workflow illustrating correlation analysis between open chromatin and gene expression and subsequent cross-referencing analysis with the EBI-GWAS data. B, Volcano plot of correlation analysis result, positively and negatively correlated relationship with p-value cutoff (0.05, one-sided Z-test). C, Genome browser view of examples of positively correlated open chromatin regions to gene expression. D-E, Correlation in chromatin openness at *VEGFA* promoter (non-significant) (D) and at *VEGFA* putative enhancer regions (significant) in relation to *VEGFA* gene activity in each cluster. F, Predicted regulatory strength of *VEGFA* putative enhancer regions estimated by a Randomforest regression model. G, Bar plots of per cent of mapped genomic locations of open chromatin regions with GWAS SNPs. H, Pie chart of per cent of classified kidney related traits to open chromatin regions. I, Genome browser views of intronic regions of *PLA2R1*/*Pla2r1* showing human-specific chromatin accessibility at LiftOver-matched genomic regions. J, Human-only, podocyte-specific *PLA2R1* regulatory region and *MAFB* binding prediction in vicinity of rs17831251 GWAS SNP locus. *See also Figure S6*

We assumed chromatin accessibility at putative enhancers correlates with gene expression levels in neighboring cell-type-specific genes, benchmarking the co-variation strategy inspired by ^71^, using cluster averaged gene expression level and open chromatin accessibility (see Methods). Adopting this co-variation approach, among the total 15,387 cell-type specific open chromatin regions, we identified 5,409 open chromatin regions (35.2%) highly correlated with the expression level of 1,468 cell-type specific genes (*p-value* < 0.05, one-sided Z-test), further listed as “putative CREs” (Table S7) (Figure 5B). The majority of highly correlated peak-to-gene pairs showed positive correlation, including putative enhancers in the previous studies (e.g., *Six1/2*^29^) For example, correlating *HNF4A* expression and open chromatin levels for each cluster identified putative CREs for the PT determinant *HNF4A*, conserved between human and mouse, corresponding to promoter-co-accessible regions and ChIP-seq verified *HNF4A* binding sites (Figure 5C). Interestingly, *HNF4A* gene activity is likely to be associated with lncRNA *HNF4A-AS1* promoter and intronic regions, suggesting a potential regulatory hierarchy between the TF and lncRNA pairs^72^.

Conversely, a minor portion of correlated gene expression-open chromatin accessibility pairs (9.74%, 681 pairs) negatively-correlated chromatin accessibility to gene expression levels (i.e., lower chromatin accessibility at cells with higher expression and highly accessible at cells with no expression), implying the importance of transcriptional repression in the precise control of transcription activity in non-lineage cells ^73^ (e.g., *SLC39A11*) (Figure S6A).

Interestingly, 294 DEGs exhibit promoter opening without obvious regulatory associated distal DARs. *VEGFA* encodes a key regulatory signal for vascular endothelial cell programs and is strongly expressed by vascular associated podocytes. *VEGFA* shows uniformly accessible (p-value: 0.39) promoter proximal DARs in all N-UE cell-type snATAC clusters (Figure 5D). However, two distal intergenic regions located at the 21.8 kb (chr6:43,744,329-43,745,600), 24.6 kb (chr6:43,746,183-43,748,431) upstream region are exclusively accessible in podocyte clusters, and their presence correlated significantly with gene expression in podocytes (*p-value*: 24.6 kb - 0.0015; 21.8 kb - 0.014) (Figure 5E). Thus, *VEGFA* expression appears accessible for activation in multiple cell types, and *VEGFA* may be transcribed at low basal levels not detected here. Elevated expression in podocytes likely reflects podocyte-specific CREs for which the distal DARs are strong candidates. We further estimated the regulatory strength of these regions by implementing a randomForest regression model per each gene and its neighboring putative CREs using the caret R package^74^. Of *VEGFA* putative CREs, the 24.6kb putative enhancer was predicted as a primary putative enhancer with the highest regulatory strength, followed by the 21.8kb distal intergenic region (Figure 5F).

To compare predicted gene regulatory insight with human disease linkage, we intersected cell-type specific DARs in the human integrated snATAC-seq clusters with all risk variant loci (NHGRI-EBI-GWAS database (v1.0.2))^75^, resulting in a total of 7,537 SNPs located within the 5,938 human DARs (Table S7). The large portion of cross-referenced open chromatin regions with GWAS SNPs were mapped to intronic and intergenic regions in the human genome (Figure 5G). Of the studies annotated in the EBI-GWAS database, we focused on 32 kidney-related traits (e.g., “glomerular filtration rate (GFR)”, “glomerulonephritis”, “proteinuria/creatinine measurement”, “acute kidney injury/chronic kidney disease (CKD)”, “nephrolithiasis”, and “diabetic nephropathy”, See method for the complete list). A total of 222 risk variants within 198 human cell-type specific DARs were associated with these kidney traits (Figure 5H) (Table S7).

We narrowed down further 74 kidney-related GWAS loci (within 62 open chromatin regions) directly related to cell-type specific gene expression and open chromatin regions in the integrated clusters (Table S7). Significant SNPs suggested disease relationships to potential gene and cell type targets, for example: *NPHS1* and *VEGFA* with podocytes; *DAB2* and *LRP2* with PT; and *IRX1*, *SIM1*, *UMOD*, and *TBX2* with DN/UE. This set included novel CRE predictions from our integrative analysis, for example: *WIPF3* with podocytes; *A4GALT* and *GATM* with PT; *TFEB* with DN; and *SPIRE2* and STC1 with UE (See Figure S6B-F).

These analyzes identified human-specific podocyte gene activity (snRNA-seq) for *PLA2R1* and its putative intronic regulatory elements (snATAC-seq), uniquely accessible in human podocytes (Figure 5I). One predicted CRE contains a highly correlated disease-risk variant associated with membranous nephropathy (MN, GWAS *p-value*: 5×10^−103^ ^76^). A predicted binding site for the podocyte regulating TF *MAFB* within 3-bp of the reported SNP suggests the SNP associated C-to-T conversion may contribute to kidney disease by altering human *PLA2R1* expression levels in human podocytes (Figure 5J). The absence of a correlated mouse *Pla2r1* expression and regulatory profile, underscores the need for appropriate human disease models. In summary, by leveraging integrated pseudo-multimodal datasets and publicly available human disease variant information, we further catalogued and prioritized CRE candidates, to consider the linkage of the human regulatory genome to human kidney disease.

## DISCUSSION

Although human and rodent species share comparable genetic blueprints reflecting conserved development and outcomes^77^, the direct extrapolation between species is becoming increasingly challenged as our understanding of human biology deepens. The high failure rate in drug development represents in part the divergence between animal model systems commonly used for drug development and divergent human biology^78^. Systemic approaches have been recently addressed to human-specific features in different organ systems or to certain specialized cell-types^79^. Single-cell profiling of the developing and adult human and mouse kidney have provided important insight within each species to developmental mechanisms and cell diversity ^10,14,21, 80–86^. By first assessing novel mouse and human multimodal datasets, then integrating mouse and human data, our study generated new insight into shared and distinct features of gene regulation and gene expression in the developing human and mouse kidney. These data and analyzes provide a valuable resource for future cross-species comparison and analysis.

Interestingly, a large fraction of genes with human-specific expression are associated with intercellular interaction, cell migration with localization to the plasma membrane. Human-specific interactions amongst kidney cells may associate with human-specific features of cell organization, enhanced organ size and extended developmental period. As an example, nephron progenitors undergo a large expansion over many weeks in a quite distinct nephrogenic niche from the mouse counterpart^21,22^. Here, human specific expression of the protocadherin *PCDH15* suggests a potential cohesive role for this Ca2+ dependent cell adhesion molecule in the human nephrogenic niche. Specifically, multiple genes were associated with neuronal development terms such as “axon guidance”. We confirmed predicted human specific expression of *NTNG1* encoding a member of the Netrin family of guidance signals in podocytes, adding new insight to published data^87,89^. Given the critical role of podocytes in harnessing the glomerular vascular endothelial network and supporting mesangial cells as a structural component, scaling between the mouse and human glomerulus may reasonably have required additional guidance cues in the human kidney.

Disease modeling using stem cell-derived organoids provides unique opportunities to characterize human diseases, enabling efficient multiplexed drug screening in the human native context^88,90^. By combining the catalogued human putative CREs, we finally focused on 62 CREs containing 74 SNPs relevant to the selected kidney-related mapped traits. Our analyses of 42 reported genes include well-studied genes with variants associated with chronic kidney disease (CKD) (Table S7), e.g., *UMOD*, *DAB2*, *LRP2*, *SLC34A1*, *SLC7A9*, and VEGFA reported^91–93^. Furthermore, TF genes linked to GWAS disease variants revealed risk alleles critical for kidney development and function. We identified human putative CREs associated with glomerular filtration rate (GFR) annotated with distal nephron program TFs including *IRX1*, *SIM1*, and *TBX2*. Furthermore, we listed 12 human congenital anomalies of kidney and urinary tract (CAKUT)-related genes (*AHI1*, *CD96*, *EVC*, *GLI2*, *GLI3*, *GREB1L*, *KANK4*, *MID1*, *PCSK5*, *ROR2*, *SEMA3A*,and *TRPC6*), by intersecting “human-specific” gene expression lists with the previously curated list in van der Ven et al., 2018 and Subramanian et al., 2019 (Table S7).

Ambiguity remains with respect to cell diversity and lineage specification for several under-represented cell-types. In the mouse nephrogenic lineage, PECs (*Cldn1^+^/Akap12^+^/Wnt16^+^,* cluster 14; Figure 3A) are predicted to arise from two distinct lineages, *Wt1*-expressing podocyte precursors and *Osr2*-expressing proximal tubule precursors: *Osr2* gene activity has been shown to label both proximal tubule precursors and maturing glomeruli in *Osr2^IresCre^*mouse^96^. In the human kidney, heterogeneity in PEC sub-types maps to flattened podocyte-like parietal cells (“parietal-podocytes”) located at the vascular pole and cuboidal *HNF4A*^+^ PECs at the tubular pole^50,97^. *Corin* expression is a specific marker of SDL in the mature adult mouse kidney ^10^. However, we observed *Corin^+^* cells co-expressing Hnf4a and *Pou3f3*+; regulatory determinants of proximal tubule and distal tubule identities, respectively^98,99^. Tracking the trajectory of *Slc12a1^+^/Foxq1^+^*TAL2 cells, RNA velocity points to a potential dual origin from either an *Slc12a1^+^/Ptger3^+^*TAL1 or *Slc12a3^+^* DCT cell lineage. Clarity on cell and lineage relationships will require the development and application of new approaches to track cell types within the developing nephron.

### Limitations of the study

We acknowledge potential pitfalls of the direct comparison in two species caused by integrating single-cell/nuclei data in distinct sequencing technologies (snRNA-seq in human versus scRNA-seq in mouse) and/or due to difficulties on matching developmental timelines (human 11.2-17.6Wk versus mouse P0). Although we independently investigate the commonality features via the step-wise analyses (Figure. 1–3), then proceeded to integrate them to curate human-specific snRNA-seq features (Figure. 4–5), we cannot exclude subtle variability on snRNA-seq and scRNA-seq differences, considering technical disparity between captured mRNA in the cell nuclei (snRNA-seq) and the whole cell (scRNA-seq)^100^.

Significant variability has been reported in mRNA transcript levels and encoded protein levels^101^, so distinct differences in mRNA levels may not translate to actual differences in functional protein contribution, and of course, many additional layers of regulation exist post-translationally that can regulate protein activity, directly or indirectly.

In the developmental survey, our in silico developmental trajectory analysis and lineage relationship is based on the RNA velocity dependent on the UMAP clustering then followed by Monocle2. Thus, the predictions in intermediate cell-types and transient gene expression should be rigorously validated by formal lineage tracing studies.

Lastly, considering kidney development involves reciprocal iterative interactions among distinct progenitors, developing interstitium and forming vascular network need to be addressed in future comparative studies. Further, given the potential local interplay of interstitial cell types in radial patterning in the kidney^106^, human-mouse differences here might have a significant impact on critical anatomical features of divergence, such as the multi-lobular architecture of the human kidney.

## Acknowledgments

The authors thank members of the McMahon laboratory and the Lindström laboratory for helpful comments and useful discussion on single cell analyses. Work in APM’s laboratory was partially supported by grants DK054364, an RBK partnership grant DK126024 and CZI grant WU-20-101 as part of the Seed Network of the Human Cell Atlas consortium (HCA).

## Author contributions

APM conceived the study. KK performed single-cell data collection. SK, RKP, MA, JS designed and/or analyzed data in consultation with APM and NOL. JG and SK performed secondary verification and followed up studies of single cell data predictions. SK assembled figures. SK, APM wrote the manuscript incorporating comments from all authors.

## DATA DEPOSITION

The single cell RNA and single nuclei ATAC sequencing data is deposited at GEO accession number: GSE232482; publicly available as of the date of publication.

## STAR Methods

### KEY RESOURCES TABLE

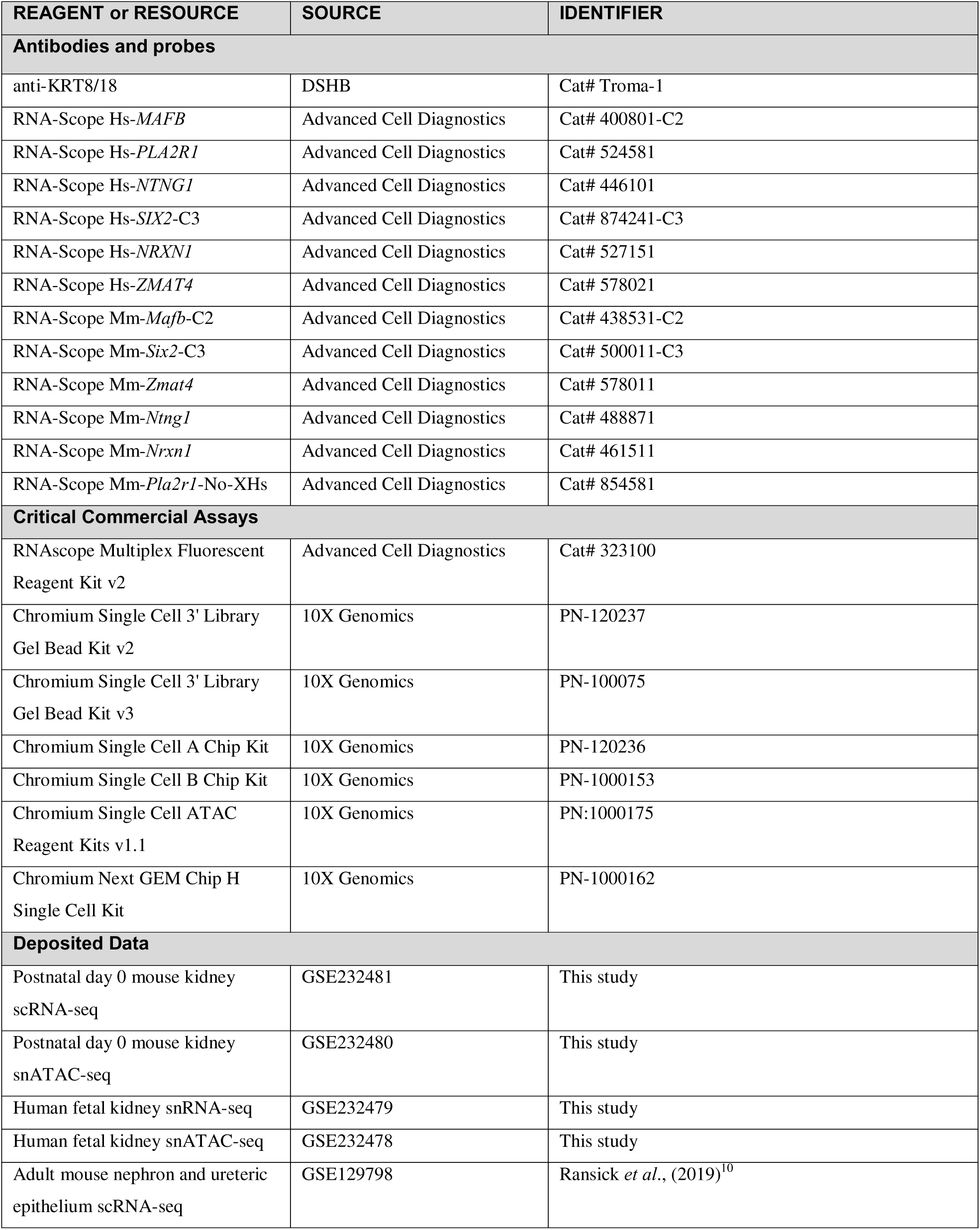
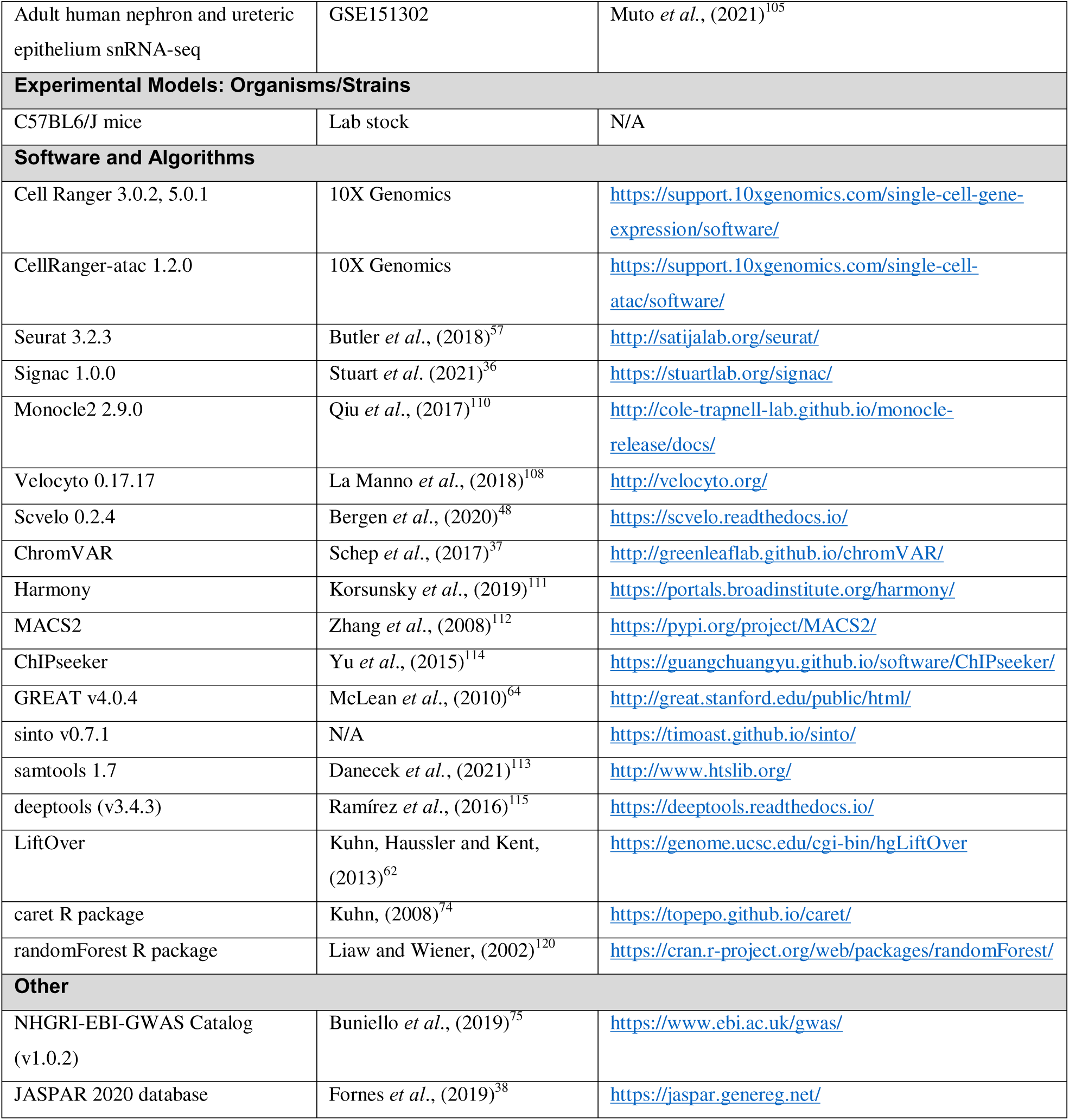

### CONTACT FOR REAGENT AND RESOURCE SHARING

Further information and requests for resources and reagents should be directed to and will be fulfilled by the Lead Contact, Andrew P. McMahon.

### EXPERIMENTAL MODEL AND SUBJECT DETAILS

Institutional Animal Care and Use Committees (IACUC) at the University of Southern California reviewed and approved all animal work as performed in this study. All work adhered to institutional guidelines. C57BL6/J mice pups were harvested at day of birth (postnatal day 0) and sexed by urogenital track (scRNA-seq; 7 males and 6 females, snATAC-seq; 2 males and 2 females). Human sample collections and analysis were performed with appropriate institutional approvals in consultation with Keck School of Medicine Institutional Review Board. Fetal embryonic age is estimated based on the first day of last menstrual period.

### METHOD DETAILS

#### Single cell preparation of P0 mouse kidney

Harvested P0 mouse kidneys were dissected into smaller pieces in cold Dulbecco’s phosphate-buffered saline (DPBS Ca2^+^-/Mg2^+^-, Gibco 14190-250). For cell isolations, the Bacillus licheniformis cold active protease (CAP) protocol was used to dissociate kidney tissue into cells as previously described in ^10^. Briefly, tissue was lysed in the DPBS lysis buffer, with the final concentration of a 2.5 mg/ml type 2 collagenase (Worthington, #LS00417), 7.5 mg/ml B. licheniformis protease (Creative Enzymes, NATE-0633), and 125 U/ml DNase I (Worthington, #LS002058) at 12°C. After 30 mins of digestion, the reaction was terminated with 20% fetal bovine serum (FBS) in DPBS. Cells were then strained through coarse 40 um cell strainers (Falcon, #352340) and pelleted at 364g in a swinging bucket centrifuge at 6°C. Cells were resuspended in cold AutoMACs Running Buffer (AMB, Miltenyi Biotec) and strained again with fine mesh 40um strainer (VWR, #76327-098). Live cells with DNA staining (DAPI-/DRAQ5^+^) were sorted by ARIA II FACS (BD Biosciences) and pelleted at 350g for 10mins then resuspended in ice cold AMB.

#### Single nuclei isolation of P0 mouse kidney

From the harvested P0 mouse kidneys, nuclei were isolated as previously described in^13^. Briefly, flash frozen kidney pieces were thawed on ice and dissected into smaller pieces (<1mm) with a razor blade and then dounced in cold Nuclei EZ Lysis Buffer (Sigma, N3408) supplemented with protease inhibitor (Roche; 05892791001) and RNAse inhibitors (Promega, N2615; Thermo Fisher Scientific, AM2696). The tissue was dounce grinded 30 times with a loose pestle (Sigma, P0485), and then filtered through a 200um strainer (pluriSelect, 43-50200). The tissue was dounce grinded 10 times with a tight pestle, and then incubated for 5 mins on ice. Cells were filtered through 40uM filter (pluriSelect, 43-50040) and pelleted at 500g for 5min at 4°C in a swinging bucket centrifuge. Supernatant was removed and nuclei pellet resuspended in Nuclei EZ lysis buffer with RNAse inhibitors, incubated an additional 5 mins on ice and pelleted again. The final pellet was resuspended in 4°C Nuclei Buffer (10X Genomics, #2000207).

#### Single nuclei isolation of human fetal kidney

Cryopreserved human fetal kidneys were processed with optimized single-nuclei dissociation protocol to minimize potential tissue damage artifacts from frozen samples and loss of intact RNA molecules during the dissociation process. Briefly, the tissues were transported in DMEM (5% FBS, 25mM HEPES), removed from capsules, dissected into <2mm pieces and flash frozen in liquid nitrogen for later isolation. Nuclei isolation from flash frozen tissue was performed by the same protocol used with mouse tissue described above. For sex identification, a small amount of the donor tissue was lysed, and DNA was isolated by incubating in DirectPCR Lysis Reagent (Qiagen, #102-T) with 1% Proteinase K (Roche, 3115879001). Donor tissue was then sexed using primers for Y-specific human SRY (SRY-FOR: CCC ATG AAC GCA TTC ATT GTG TGG/ SRY-REV: ATT TTA GCC TTC CGA CGA GGT CGA TA), and ZFX/ZFY (ZFX/ZFY FOR: GCA CTT CTT TGG TAT CTG AGA AAG T/ ZFX/ZFY REV: ATA ATC ACA TGG AGA GCC ACA AGC T) to confirm DNA isolation quality.

#### sc/snRNA-seq and snATAC-seq library construction and sequencing

The sequencing library information used in this study is summarized in Supplementary table #. For sc/snRNA-seq, Chromium Single Cell 3’ Reagent Kits v2/3 reagents kits (10X Genomics, v2; PN-120237, v3; PN-100075) were used to generate single cell 3’ gene expression libraries according to manufacturer’s instructions (10X genomics, v2; CG00052, v3; CG000183). Briefly, freshly prepared single-cell/nuclei suspension in the corresponding resuspension buffers was combined with 10X Chromium reagent master mix. Target 8,000 cells (mouse) or 4,000-6,000 nuclei (human) ascertained by Trypan Blue hemocytometer count were loaded into 10X Chromium Chips (v2; Chip A, v3; Chip B) for GEM generation, barcoding, and library construction. All thirteen libraries for mouse scRNA-seq and two human snRNA-seq libraries were sequenced in HiSeq4000 platform (Illumina) at Translational Genomics Center at Children’s Hospital Los Angeles Center (CHLA) for Personalized Medicine. Among eleven human snRNA-seq libraries, four were sequenced on DNBseq-G400 platform at BGI Genomics and five were sequenced on Illumina-S4 Novaseq 6000 platform at Novogene.

For snATAC-seq, Chromium Single Cell Reagent Kits v1.1 (10X Genomics, PN-1000175) reagents and index plate were used to generate the snATAC-seq libraries per the manufacturer’s specifications (10X Genomics, CG000209). Briefly, target 10,000 mouse or human nuclei ascertained by Trypan Blue hemocytometer count were proceeded to Tn5 transposition then loaded into 10X Chromium Chips (Chip H, 10X Genomics, PN-1000162) for GEM generation, barcoding, and library construction. All four mouse snATAC-seq libraries and four of nine human snATAC-seq libraries were sequenced on DNBseq-G400 platform at BGI Genomics. Five of nine human snATAC-seq libraries were sequenced on Illumina-S4 Novaseq 6000 platform at Novogene. (Table S2)

#### In Situ Hybridization (ISH) and Immunofluorescence (IF) Co-detection Analyses

We modified the RNAscope® Multiplex Fluorescent Reagent Kit v2 (Advanced Cell Diagnostics) to include immunofluorescent staining procedures for co-detection of target mRNAs and proteins. Briefly, neonatal (P0) mouse kidney and human fetal (17.2 week) cryosection slides were washed in PBS and dried at room temperature for 10 minutes. The slides were treated with hydrogen peroxide, protease 3, then hybridized with RNAScope probes for 2 hours in 40’C using the HybEZ oven as recommended. The RNAscope probes from Advanced Cell Diagnostics used in this work are listed as follow: Hs-*MAFB* (400801-C2), Hs-*PLA2R1* (524581), Hs-*NTNG1*(446101), Hs-*SIX2*-C3 (874241-C3), Hs-*NRXN1* (527151) Hs-*ZMAT4* (578021), Mm-*Mafb*-C2 (438531-C2), Mm-*Six2*-C3 (500011-C3), Mm-*Zmat4* (578011), Mm- *Ntng1*(488871), Mm-*Nrxn1* (461511), and Mm-*Pla2r1*-No-XHs (854581). The hybridization signals were amplified and developed with TSA plus fluorophores. To incorporate immunostaining, the slides were incubated with the blocking solution, 1.5% Seablock (ThermoFisher) and 0.25% TritonX-100 in PBS, for 1 hour at room temperature, then incubated with primary antibody KRT8/18 (DSHB, troma-1; 1:50) in 4’C, overnight. Alexa Flour-conjugated secondary antibodies diluted in blocking solution were applied for 1 hour at room temperature, and then nuclei counterstaining was performed using 1 µg/ml Hoechst 33342 (Molecular Probes) in PBS. The slides were mounted with Prolong Gold Antifade Reagent (Life technologies) then imaged at either 20X or 40X using the Leica Sp8 confocal microscope. The images were processed and exported by Leica Application Suite X (LAS X) Microscope software (Leica) into Adobe Illustrator (Adobe). Complete details of antibodies and RNAscope probes in tabular form in Key Resources Table.

### QUANTIFICATION AND STATISTICAL ANALYSIS

#### Single cell RNASeq Data Analysis

##### Preprocessing, quality-control, dimension reduction, dataset merge/integration

All sc/snRNA-seq data was processed via the Cell Ranger Single-Cell Software (mouse scRNA-seq; v.3.0.2, human snRNA-seq; v.5.0.1) and Seurat R package (v.3.2.3). Briefly, sequencing raw FASTQ files were processed by the Cell Ranger ‘cellranger count’ command to align to the pre-built references for mouse (mm10-3.0.0) and human genomes (GRCh38-2020-A). We used ‘include-introns’ option in snRNA-seq in the Cell Ranger alignment to ensure a higher number of genes detected per nucleus and better cell-type identification. We reviewed the alignment summary reports and excluded one replicate of the human snRNA-seq libraries due to its poor library construction and alignment quality metric. The filtered feature-barcode matrices were loaded into Seurat with Read10X function.

For mouse scRNA-seq data, seven samples (10X Chromium Chemistry v2) and six samples (10X Chromium Chemistry v3) for mouse scRNA-seq datasets were merged into two individual Seurat objects according to the matching reagent versions and normalized individually with SCTransform^102^ to regress out mRNA transcript per cell. The two Seurat objects were applied quality-control metrics for genes per cell (500-5,500), mRNA transcripts per cell (500-30,000), maximum mitochondrial percentage (35%) to eliminate low quality cells. To minimize potential batch effects caused by the distinct library preparation reagents, these objects were integrated into a single unified mouse scRNA-seq data object with all thirteen mouse replicates via the CCA-based integration implemented in Seurat V3, through the standard Seurat data integration pipeline on datasets normalized with SCTransform. After removing weakly clustered cells, non-linear dimension reduction was performed via RunUMAP and FindNeighbors function with 30 principal components (PCs) from ‘integration’ assay layer, followed by graph-based clustering using FindNeighbors and FindClusters at resolution 1.0.

For human snRNA-seq data, six samples sequenced on DNBSEQ-G400 and HiSeq4000 and four samples on Illumina-S4 Novoseq6000 were individually merged, normalized, quality control-filtered, then integrated to a single human dataset as described above in the mouse scRNA-seq data processing. Weakly clustered cells with poor quality metrics were iteratively removed in the two rounds of dimension reduction with the same parameters used in the mouse scRNA-seq described above.

#### Differential gene expression analysis, cluster identification, and subclustering analysis

Differential gene expression analysis was performed based on normalized transcripts per cell in the ‘RNA’ assay layer via FindAllMarkers function with the default parameters to identify positive markers for each cluster. Clusters are identified based on known cell-type specific markers and further annotated according to the top differentially expressed genes and the UMAP clustering. Selected cluster numbers for different cellular groupings – 1) N-UE, 2) interstitial, 3) vascular, and 4) immune cell-types – were separated with subset function. As ‘integration’ assay layers were computed during the first integration step previously, subsetted cells with different reagent versions or sequencing platforms are individually treated to re-compute variable features for each grouping. These variable features were used to re-integrate and cluster cells by using 20 PCs. These objects are used for the downstream subclustering analysis without more iterative integration processes.

#### TF-focused analysis in human and mouse N-UE transcriptomic data

To assess the specificity and diversity of transcriptional regulators expressed in the human and mouse N and UE lineage cells in the sc/snRNA-seq datasets, we examined the gene expression levels of transcription factors listed in the previous mouse^103^ and human^104^ TF atlases. Transcription factors that were expressed by 5 or more cells of each dataset were selected and their average expression levels were calculated by AverageExpression function in Seurat (Table S3). The specificity of representative conserved mouse or human orthologs in Ensembl BioMart ^107^ was visualized as the dotplots in Figure S2.

#### Correlation analysis with the published human and mouse adult scRNA-seq datasets

The adult human and mouse kidney transcriptome datasets (mouse adult kidney^10^, human adult kidney^105^) and were downloaded and loaded into Seurat. Averaged gene expression matrices per cluster were computed by using AverageExpression function. Gene expression levels in the commonly expressed and variable genes were listed with VariableFeatures and intersect functions. Spearman correlation was computed by cor function under the consideration of rank-ordered variables in distinct datasets. Pearson correlation matrices were converted into heatmap representations by pheatmap (v1.0.12) R package.

#### Generating nephron-only compartments and lineage assignment

To analyze the human and mouse nephrogenic lineage cells in the finer detail, the cells within the nephron (N) compartment were computationally selected by the UMAP clusters. By the Hox gene profiling, *HOXD/Hoxd10/11*^+^ NPC-derived collecting duct cells were distinguished, separated, and re-integrated to generate more inclusive human and mouse N-lineage compartments. To avoid potential artifacts in RNAVelocity and downstream developmental analysis in the nephrogenic compartments, cells encompassing from *PAX8/Pax8*- expressing NPC to terminal cell-types were subsetted, subclustered, and refined iteratively with multiple steps eliminating proliferative NPCs. Lastly, three main nephrogenic lineages were assigned based on expression of the key transcription regulator markers (*MAFB/Mafb*, *HNF4A/Hnf4a*, and *POU3F3/Pou3f3*, for podocyte, proximal tubule cells and distal nephrons, respectively). For the *MAFB/Mafb+* lineage compartments, two terminal cell-types were identified, and parietal epithelial cell-type clusters were further excluded to generate the podocyte-specific data.

#### RNA velocity and pseudotime analysis

We utilized Velocyto (version 0.17.17, ^108^) and Scvelo (0.2.4, ^48^) for RNAVelocity analysis to predict developmental relationship from progenitor cell-types to terminally differentiated cell-types. Read counts in BAM files from individual CellRanger alignment output folders were imported as loom files by velocyto run, loaded by anndata read_loom ^109^, and merged into a single combined file using loompy connect. Additional metadata information in target Seurat object, including cell barcodes, UMAP cluster numbers, and UMAP dimensions, were exported from target Seurat objects and transferred into loom files. Follow by preprocessing with scv.pp.filter_and_normalize, velocity estimation was performed by scv.pp.moments and scv.tl.velocity with the “stochastic” mode.

We inferred lineage specification processes by examining Velocity arrow transitions and known developmental and terminally differentiation markers, then computed pseudotime analysis using Monocle2 (2.9.0, ^110^). Additional subclustering was performed if necessary for greater details. Seurat objects are converted into Monocle2-compatible CellDataSet objects by as.CellData. Pseudotime ordering was calculated by reduceDimension function with method parameter ‘DDRTree’ and orderCells function, based on the top 10% differentially expressed genes in target Seurat objects.

#### Single nucleus ATACSeq Data Analysis

##### Preprocessing, quality-control, dimension reduction, dataset merge/integration

All snATAC-seq data was processed via the Cell Ranger ATAC Software (v.1.2.0) and Signac (v.1.0.0) R package as an extension for Seurat. Briefly, sequencing raw snATAC-seq data was aligned to the pre-built references for mouse (atac-mm10-1.2.0) and human genomes (atac-GRCh38-1.2.0) followed by peak-calling with ‘cellranger-atac count’ command, to generate BAM files, called peak files, filtered peak-barcode matrices, and barcoded and aligned fragment files. Called peak BED format files and fragment files were loaded into Signac with read.table function and CreateFragmentObject function.

For mouse snATAC-seq data, an initial filtering of cells with low fragment counts (<500) was performed for the four mouse snATAC-seq libraries. Genomics ranges of peaks in the datasets were intersected to obtain a set of overlapping genomic regions over all datasets with subsetByOverLaps function. Peaks then were primarily filtered with the range of peak width 20-10,000bp. Each replicate was further individually filtered with quality control metrics (peak_region_fragment 2,000-50,000; pct_reads_in_peaks > 30; blacklist_ratio < 0.05, nucleosome_signal < 2; TSS.enrichment > 2), then merged into a single dataset with merge function. Based on the intersected common peak set, normalization and linear dimension reduction was performed with the latent semantic indexing (LSI) implemented in Signac, which is the combination of TF-IDF (term frequency-inverse document frequency) normalization, feature selection for peaks with more than 50 nuclei, and singular value decomposition (SVD). Graph-based clustering and non-linear dimension reduction was performed with the first 30 components except for the first LSI component with FindNeighbors, FindClusters, and RunUMAP.

“Gene activity” score was calculated to assess the chromatin accessibility associated with each gene, summing total ATAC fragments fall into gene bodies and 2kb upstream regions of transcription start site (TSS) according to the Bioconductor R package EnsDb.Mmusculus.v79.

Motif information based on position frequency matrices (PFM) fetched from JASPAR 2020 database^38^ was added on the commonly intersected peaks. Base composition information was used to suppress overrepresented motifs by normalizing the GC content, region lengths, and dinucleotide base frequencies with RegionStats function based on with the Bioconductor R package BSgenome.Mmusculus.UCSC.mm10 reference genome. Per nuclei motif activity scores were computed via ChromVAR^37^ with RunChromVAR function.

For human snATAC-seq data, four samples sequenced on DNBSEQ-G400 and five samples on Illumina-S4 were individually merged into two Signac objects and applied with quality metrics (peak_region_fragment 2,000 - 50,000; pct_reads_in_peaks > 30; blacklist_ratio < 0.05, nucleosome_signal < 2; TSS.enrichment > 2) as described in the mouse snATAC-seq datasets. Harmony integration^111^ was performed to integrate the two filtered object into a single dataset, based on the intersected peak regions within the two objects obtained by findOverlaps and queryHits functions. By using RunHarmony function, returned batch effect-corrected LSIs were used for dimension reduction and clustering. Gene activity score and motif information were added to the Signac objects as described in the mouse snATAC-seq data, according to the Bioconductor R packages EnsDb.Hsapiens.v86 and BSgenome.Hsapiens.UCSC.hg38 reference genome.

#### Differential accessible region, gene activity, and subclustering analysis

Differential accessible region (DAR), gene activity (GA) and motif activity analysis were all performed by FindAllMarkers function. For DAR and motif activity, differential analysis utilized the logistic regression (LR) with a latent variable of the total number of fragments in peak regions per nuclei on the respective layers. For GA, differential analysis was performed with the default parameters. Subclustering analysis was done as described in the sn/snRNA-seq section, then another round of peak calling by using CallPeaks function, with MACS2^112^ backend, was used to list differentially accessible regions at the final subclustering clusters.

#### Genomic region and regulatory domain annotation of cell-type specific open chromatin regions

Differentially accessible regions calculated in snATAC-seq data were further annotated to have functional interpretation and regulatory context. We utilized ChIPseeker annotatePeak function with default parameter^114^ for genomic region annotation of each open chromatin region (e.g., Promoter, 5’ UTR, 3’ UTR, Exon, Intron, Downstream, and Intergenic). Transcript-related features and reference genome information of mouse reference (mm10) or human reference (hg38) genome were obtained from Bioconductor R package TxDb.Mmusculus.UCSC.mm10.knownGene, org.Mm.eg.db (mouse), TxDb.Hsapiens.UCSC.hg38.knownGene, and org.Hs.eg.db (human). To harness genomics regions to genes as regulatory domains, GREAT v4.0.4^64^ is used to annotate open chromatin regions with more than average fold change 0.5 (based on Seurat FindAllMarkers result, avg_FC > 0.5) to their associated genes with regulatory domain parameter “Basal plus extension”.

#### Visualization of single-cell/nuclei data

Most single-cell/nuclei plots were visualized by the Seurat or Signac plotting functions guided by the visualization vignettes, using UMAPPlot, DotPlot, FeaturePlot functions. For aggregated pseudo-bulk ATAC-seq plots in a genome browser-like view, cell/nucleus barcodes in individual clusters were specified with WhichCells function. BAM files per sample generated from the Cell Ranger ATAC software outputs were split into different clusters by filterbarcodes command from the Python package sinto (v0.7.1, https://github.com/timoast/sinto) then merged again with samtools (v1.7)^113^ merge command. BAM files per cluster were converted into BigWig files with bamCoverage command from deeptools (v3.4.3)^115^ (--binSize 100 -- ignoreForNormalization chrX --normalizeUsing RPGC –extendReads) and loaded into IGV (v2.8.7) browser or UCSC genome browser for visualization.

#### Integration of human and mouse transcriptomic and open chromatin accessibility atlases

To integrate human and mouse N-UE sc/snRNA-seq and snATAC-seq datasets into a single unified atlas, we leveraged canonical correlation analysis (CCA) integration implemented Seurat V3^57^ in a step-wise manner. First, human and mouse N-UE sc/snRNA-seq cells were integrated on the basis of common orthologous variable features. Briefly, mouse gene symbols in mouse scRNA-seq Seurat objects were renamed to human orthologous gene symbols ^116^. The CCA-based integration was performed with 3,000 most variable features selected by SelectIntegrationFeatures function, then cross-dataset pairs of cells in matched cell-types, i.e., ‘anchors’, were identified FindIntegrationAnchors function. Cross-species human-mouse sc/snRNA-seq data were generated though IntegrateData function and applied to another round of clustering to define equivalent, corresponding human and mouse cell-types. Second, by using cell-type clustering information in the cross-species sc/snRNA-seq data as a reference atlas, the human and mouse snATAC-seq datasets were subsequently incorporated through label transferring and data imputation as described in the Seurat V3 co-embedding workflow. Pseudo-multimodal integration anchors were identified via FindTransferAnchors between variable gene expression features (sc/snRNA-seq) and variable gene activity features (snATAC-seq), then RNA-to-ATAC data imputation was done by TransferData function. As the human and mouse sc/snRNA-seq integration yielded shared data dimension metrics in clustering, the co-embedded human and mouse RNA-ATAC-seq datasets were merged and re-clustered into a single UMAP landscape with the inclusive alignment of different species and modality layers.

#### Conserved and species-specific molecular features in the integrated cross-species pseudo-modal dataset

We identified commonly expressed cell-type DEGs between corresponding human and mouse N-UE cell-types by using FindConservedMarkers function with group variable “species”, listed 1,390 conserved genes. To have more cell-type specificity, 352 genes were listed after applying applied another filter with higher fold changes (avg_logFC > 0.5) and per cent of expressing cells (pct.1 – pct.2 > 0.3). Species-specific differentially expressed genes are listed by excluding conserved DEGs in human and mouse data from one another, then used the same criteria with conserved gene. For conserved and divergent DARs, we converted human or mouse DARs to mouse or human reference genome (hg38→mm10, or mm10→ hg38) based on aligned syntenic regions (“UCSC chain files”), by using LiftOver tool^62^, then overlapping regions between human and mouse regions by using findOverlaps function in IRanges R package^117^. Conserved and species-specific DARs were identified based on whether LiftOver-converted regions are corresponding each other or not matching/no syntenic regions found.

#### Peak-gene cis-association by covariation analysis and regulatory potential of correlated peaks

In the pseudo-multimodal integration, each integrated cluster contains cell-type cluster-matching differentially expressed genes and differentially accessible regions. We adopt and modified the peak-gene cis-association computational method developed in the Share-seq paper ^118^. For simplicity, we targeted genes and their open chromatin regions that were annotated by GREAT annotation described above (regulatory domain parameter, “Basal plus extension”). To calculate peak-to-gene association and regulatory relationship, we used gene expression values and associated open chromatin region peak counts from each cluster to calculate the observed Pearson correlation (obs). We calculate the Pearson correlation of randomly selected 100 GC content-matching background open chromatin regions with gene expression values, to normalize potential biases of GC percentages. The Pearson correlation (pop.mean) and its standard deviation (pop.sd) were used for one-sided Z test (z = (obs-pop.mean)/pop.sd)^118^ to determine p-values. We defined highly-correlated peaks (p-value < 0.05) as “putative cREs”. To examine putative cCREs and co-accessible regulatory domains, co-accessibility analysis by Cicero^119^ was performed to generate cis-regulatory networks between promoter regions and GREAT-annotated neighboring peaks. To determine relative regulatory potential of each peak to gene expression levels, we constructed Randomforest regression model with openness of putative cREs (input/features) predicting imputated gene expression levels (output/labels) per cells with caret^74^ and randomForest^120^ R package (method=’rf’, tuneLength=25) with repeated 10-fold cross validation. Feature variable importance of each peak was examined by varImp function with randomForest model outputs.

#### Incorporating kidney health trait and disease context to cell-type specific gene activity and open chromatin accessibility

To incorporate risk alleles to kidney health traits and diseases, SNP genomic loci from the publicly available GWAS catalog (NHGRI-EBI-GWAS database (v1.0.2), ^75^) were intersected with human cell-type specific DARs. A tabulated list of SNPs was cross-referenced by using findOverlaps function from IRanges R package. Total of 32 kidney-related terms (“Acute kidney injury”, “albuminuria”, “chronic kidney disease”, “creatinine clearance measurement”, “creatinine measurement”, “diabetic nephropathy”, “GFR change measurement”, “glomerular filtration rate”, “Hematuria”, “IGA glomerulonephritis”, “inosine measurement”, “kidney transplant”, “lower urinary tract symptom”, “lupus nephritis”, “membranous glomerulonephritis”, “Moderate albuminuria”, “Nephroblastoma”, “nephrolithiasis”, “nephrotic syndrome”, “Proteinuria”, “renal cell carcinoma”, “renal system measurement”, “renal transplant outcome measurement”, “susceptibility to urinary tract infection measurement”, “type 1 diabetes nephropathy”, “urinary albumin to creatinine ratio”, “urinary metabolite measurement”, “urinary microalbumin measurement”, “urinary nitrogen measurement”, “urinary pH measurement”, “Urinary retention”, and “urinary uromodulin measurement”) were classified into glomerular filtration rate (GFR), glomerulonephritis, proteinuria and creatinine measurement, acute kidney injury and chronic kidney disease (CKD), nephrolithiasis (kidney stones), and diabetic nephropathy. The intersected SNP loci were individually inspected in aggregated pseudo-bulk snATAC-seq tracks in conjunction to “Cons 100 Verts by PhastCons”, “ENCODE cCRE tracks” and “JASPAR2020 TFBS hg38” tracks available in UCSC genome browser. For potential association with human genetic disorders, human fetal N-UE cell-type specific DEGs were cross-referenced with Congenital anomalies of the kidneys and urinary tract (CAKUT)-related gene list from ^121^ and ^95^.

## SUPPLEMENTAL INFORMATION

Figure S1 Dataset metrics and rare neuronal precursor-like cell-types in the human and mouse developing kidney sn/scRNA-seq and snATAC-seq Related to Figure 1

A-B, dataset as violin plots of clusters in Figure 1B-C, showing per cluster levels for nGenes (top), nUMI (middle) and percent mitochondrial genes (bottom) (B) mouse scRNA-seq (C) human snRNA-seq. C-D, dataset as violin plots of clusters in Figure 1F-G, showing per cluster levels for nUMIs (1^st^ row), nGene (2^nd^ row), nFragment in regions (3^rd^ row), fragment ratio in blacklist (4^th^ row) (C) mouse snATAC-seq (D) human snATAC-seq. E, a rare *Sox10*^+^/*Foxd3*^+^ cell-type cluster in mouse scRNA-seq, suggestive of Schwann cell precursors. F, a rare *PHOX2B*^+^/*HAND2*^+^ cell-type cluster in human snATAC-seq, suggestive of autonomic neuron precursors of neural crest derivatives.

Figure S2 The broad concordance mouse P0 and human fetal kidney cell-types and less represented mature loop of Henle sub-types Related to Figure 2

A, Pearson correlation analysis between mouse P0 (vertical) and human fetal (horizontal) cell-types showing similarity in transcriptomic features for each cluster. B-C, Pearson correlation heatmaps showing transcriptomic similarity in selected mouse P0 clusters and previously reported mouse adult^10^ clusters (B), selected human fetal clusters and previously reported human adult^105^ clusters (C) D, Harmonized view of cell-type specific TF gene activity dot plots, cluster numbers can be found on the top UMAP plots. E-G, UMAP projections of gene expression markers for cell-type diversity of loop of Henle reported in adult mouse^10^, two representative markers for SDL (E), tDL (F), and tAL cells (G).

Figure S3 Developmental trajectory analysis in the mouse nephrogenic lineage cells and cross-checking the human counterparts revealed the broad commonality and divergent details Related to Figure 3

A, RNA velocity analysis on the UMAP projections of the human and mouse N-UE cells. B, in silico separation of nephrogenic cells, Dot plot of Hox gene profiling (left) UMAP projection showing *HOXD10/11/Hoxd10/11* gene activity (right) C, Genome browser plot showing a human-specific open chromatin region at OLFM3/Olfm3. D, Pseudotime ordered mouse PT trajectory. E, UMAP projection showing Corin and *Satb2* gene activity with overlapping *Irx1* and *Pou3f3* activity in the mouse PT lineage compartment. F, UMAP projections showing gene activity of mouse cortical SDL (left; *Slc6a12*/*Bcl6*) and juxtamedullary tDL markers (right; *Aqp1*/*Pitx2*) in Bst^+^ DTL cells (cluster 13). G-J, Pseudotime ordered-individual mouse DN trajectories, DTL (G), TAL (H), DCT (I), and CNT (J).

Figure S4 Cross-species pseudo-multimodal integration and human-specific cell-type specific gene activity Related to Figure 4

A, Stacked bar plots showing per cent of cells/nuclei in clusters in integrated dataset, clusters with less or no snATAC-seq are marked by asterisks on the top. B, Harmonized view of gene activity profiles from snATAC-seq viewpoint in human/mouse integrated clusters. C, A panel of UMAP projections of *ZNF730* and *ZNF385D* encoding human zinc finger proteins with no known mouse orthologs. D, RNA scope in situ hybridization validation of human-specific gene activity of *ZMAT4*. E. A panel of UMAP projections of two TF, *NR3C2*, *NPAS2*, showing human-specific gene activity. F, A panel of UMAP projections and violin plots of genes that have different local enrichment in human and mouse (*UNC5B*/*Unc5b*, *EYA4*/*Eya4*, *FGF1*/*Fgf1*, and *SPINK1*/*Spink1*) G, Bar plots of Gene Ontology analysis of conserved and human-specific DEG EnrichR enrichment.

Figure S5 Cross-species pseudo-multimodal integration and human-specific cell-type specific open chromatin regions Related to Figure 4

A, human-specific open chromatin regions associated with human-specific gene activity; UMAP projection showing *DLG2*/*Dlg2* expression and a genome browser view at human-specific open chromatin regions. B-C, human-specific open chromatin regions associated with conserved gene activity; UMAP projection showing gene activity of *ZBTB7C*/*Zbtb7c* (B) and *CLIC5*/*Clic5* (C) and a genome browser view of human-specific open chromatin regions.

Figure S6 Human open chromatin regions highly correlated to gene expression levels and putative cis-regulatory regions potentially associated with kidney diseases Related to Figure 5

A, Genome browser views and UMAP projections showing open chromatin accessibility and relevant gene expression levels: examples of negatively correlated open chromatin regions to gene expression (A), B-F, disease-related SNPs associated with genes showing human-specific gene activity and open chromatin regions in the corresponding cell-types in the integrated dataset, *WIPF3* (B) *A4GALT* (C), *GATM* (D), *TFEB* (E), *SPIRE2* (F), *STC1* (G)

## SUPPLEMENTAL TABLES

Table S1 Complete list of human and mouse developmental sc/snRNA-seq and snATAC-seq sequencing data used in this study.

Table S2 Human and mouse global and N-UE subclustering sc/snRNA-seq and snATAC-seq data lists of differentially expressed genes (RNA) and differentially accessible regions/differential TF binding predictions (ATAC) for each cluster shown in the UMAP projections in Figure 1B-C, 1F-G, 2B-C, and 2K-O. See tab entitled “cell-type annotation” for key to content, Excel file. *Related to Figure 1, 2 and Figure S1,2*

Table S3 Human and mouse *PAX8*/*Pax8*-expressing differentiating nephrons and three lineage specification trajectory data list of differentially expressed genes shown in the UMAP projections in Figure 3B. See tab entitled “lineage assignment and terminal cell-type annotation” for key to content, Excel file. *Related to Figure 3 and Figure S3*

Table S4 Cross-species pseudo-multimodal integrated N-UE datasets list of differentially expressed genes (RNA) and differentially accessible regions/differential TF binding predictions (ATAC) for each cluster shown in the UMAP projections in Figure 4A, See tab entitled “cell-type annotation” for key to content, Excel file. *Related to Figure 4 and Figure S4*

Table S5 Conserved and human-specific differentially expressed genes and differentially accessible regions for each corresponding human/mouse sc/snRNA-seq or snATAC-seq clusters in the integration analysis in Figure 4. See tab entitled “read me” for key to content, Excel file. *Related to Figure 4 and Figure S4*

Table S6 Genes with distinct local enrichment patterns in the integration analysis in Figure 4. *Related to Figure 4 and Figure S4D*

Table S7 Highly correlated open chromatin regions to gene expression and cross-reference analysis with the EBI GWAS SNP data in the human N and UE cell-type clusters of the integrated cross-species pseudo-multimodal datasets in Figure 5. *Related to Figure 5 and Figure S5*

## REFERENCES

1. Nielsen, S., Kwan, T.-H., Fenton, R.A., and Praetorious, J. (2012). Anatomy of the kidney. Brenner and Rector’s the Kidney, 31–93.

2. Mandel, E.I., Bernacki, R.E., and Block, S.D. (2017). Serious illness conversations in ESRD. Clin. J. Am. Soc. Nephrol. 12, 854–863. 10.2215/CJN.05760516.

3. Oxburgh, L., Carroll, T.J., Cleaver, O., Gossett, D.R., Hoshizaki, D.K., Hubbell, J.A., Humphreys, B.D., Jain, S., Jensen, J., Kaplan, D.L., et al. (2017). (Re) Building a kidney. J. Am. Soc. Nephrol. 28, 1370–1378. 10.1681/ASN.2016101077.

4. McMahon, A.P. (2016). Development of the Mammalian Kidney. In Current Topics in Developmental Biology (Academic Press), pp. 31–64. 10.1016/bs.ctdb.2015.10.010.

5. Little, M.H. (2015). Improving our resolution of kidney morphogenesis across time and space. Curr. Opin. Genet. Dev. 32, 135–143. 10.1016/j.gde.2015.03.001.

6. Merlet-Bénichou, C., Gilbert, T., Vilar, J., Moreau, E., Freund, N., and Lelièvre-Pégorier, M. (1999). Nephron number: variability is the rule. Causes and consequences. Lab. Invest. 79, 515–527.

7. Hughson, M., Farris, A.B. 3rd, Douglas-Denton, R., Hoy, W.E., and Bertram, J.F. (2003). Glomerular number and size in autopsy kidneys: the relationship to birth weight. Kidney Int. 63, 2113–2122. 10.1046/j.1523-1755.2003.00018.x.

8. Oliver, J., and MacDowell, M. (1968). Nephrons and Kidneys. A Quantitative Study of Developmental and Evolutionary Mammalian Renal Architectonics… Based on the Microdissections of Muriel MacDowell. 27 Plates and 25 Text Figures (New York).

9. Dantzler, W.H., Pannabecker, T.L., Layton, A.T., and Layton, H.E. (2011). Urine concentrating mechanism in the inner medulla of the mammalian kidney: role of three-dimensional architecture. Acta Physiol. (Oxf). 202, 361–378. 10.1111/j.1748-1716.2010.02214.x.

10. Ransick, A., Lindström, N.O., Liu, J., Zhu, Q., Guo, J.J., Alvarado, G.F., Kim, A.D., Black, H.G., Kim, J., and McMahon, A.P. (2019). Single-Cell Profiling Reveals Sex, Lineage, and Regional Diversity in the Mouse Kidney. Dev. Cell 51, 399–413.e7. 10.1016/j.devcel.2019.10.005.

11. Park, J., Shrestha, R., Qiu, C., Kondo, A., Huang, S., Werth, M., Li, M., Barasch, J., and Suszták, K. (2018). Single-cell transcriptomics of the mouse kidney reveals potential cellular targets of kidney disease. Science (80-.). 360, 758–763. 10.1126/science.aar2131.

12. Chen, L., Chou, C.-L., and Knepper, M.A. (2021). Targeted Single-Cell RNA-seq Identifies Minority Cell Types of Kidney Distal Nephron. J. Am. Soc. Nephrol. 32, 886 LP–896. 10.1681/ASN.2020101407.

13. Wilson, P.C., Wu, H., Kirita, Y., Uchimura, K., Ledru, N., Rennke, H.G., Welling, P.A., Waikar, S.S., and Humphreys, B.D. (2019). The single-cell transcriptomic landscape of early human diabetic nephropathy. Proc. Natl. Acad. Sci. U. S. A. 116, 19619–19625. 10.1073/pnas.1908706116.

14. Liao, J., Yu, Z., Chen, Y., Bao, M., Zou, C., Zhang, H., Liu, D., Li, T., Zhang, Q., Li, J., et al. (2020). Single-cell RNA sequencing of human kidney. Sci. Data 7, 1–9. 10.1038/s41597-019-0351-8.

15. Young, M.D., Mitchell, T.J., Vieira Braga, F.A., Tran, M.G.B., Stewart, B.J., Ferdinand, J.R., Collord, G., Botting, R.A., Popescu, D.M., Loudon, K.W., et al. (2018). Single-cell transcriptomes from human kidneys reveal the cellular identity of renal tumors. Science (80-.). 361, 594–599. 10.1126/science.aat1699.

16. Lake, B.B., Chen, S., Hoshi, M., Plongthongkum, N., Salamon, D., Knoten, A., Vijayan, A., Venkatesh, R., Kim, E.H., Gao, D., et al. (2019). A single-nucleus RNA-sequencing pipeline to decipher the molecular anatomy and pathophysiology of human kidneys. Nat. Commun. 10, 1–15. 10.1038/s41467-019-10861-2.

17. Kobayashi, A., Valerius, M.T., Mugford, J.W., Carroll, T.J., Self, M., Oliver, G., and McMahon, A.P. (2008). Six2 Defines and Regulates a Multipotent Self-Renewing Nephron Progenitor Population throughout Mammalian Kidney Development. Cell Stem Cell 3, 169–181. 10.1016/J.STEM.2008.05.020.

18. Humphreys, B.D., Lin, S.L., Kobayashi, A., Hudson, T.E., Nowlin, B.T., Bonventre, J. V., Valerius, M.T., McMahon, A.P., and Duffield, J.S. (2010). Fate tracing reveals the pericyte and not epithelial origin of myofibroblasts in kidney fibrosis. Am. J. Pathol. 176, 85–97. 10.2353/ajpath.2010.090517.

19. Ly, J.P., Onay, T., and Quaggin, S.E. (2011). Mouse models to study kidney development, function and disease. Curr. Opin. Nephrol. Hypertens. 20, 382–390. 10.1097/MNH.0B013E328347CD4A.

20. Appel, D., Kershaw, D.B., Smeets, B., Yuan, G., Fuss, A., Frye, B., Elger, M., Kriz, W., Floege, J., and Moeller, M.J. (2009). Recruitment of podocytes from glomerular parietal epithelial cells. J. Am. Soc. Nephrol. 20, 333–343. 10.1681/ASN.2008070795.

21. Lindström, N.O., Guo, J., Kim, A.D., Tran, T., Guo, Q., De Sena Brandine, G., Ransick, A., Parvez, R.K., Thornton, M.E., Basking, L., et al. (2018). Conserved and divergent features of mesenchymal progenitor cell types within the cortical nephrogenic niche of the human and mouse kidney. J. Am. Soc. Nephrol. 29, 806–824. 10.1681/ASN.2017080890.

22. Lindström, N.O., McMahon, J.A., Guo, J., Tran, T., Guo, Q., Rutledge, E., Parvez, R.K., Saribekyan, G., Schuler, R.E., Liao, C., et al. (2018). Conserved and divergent features of human and mouse kidney organogenesis. J. Am. Soc. Nephrol. 29, 785–805. 10.1681/ASN.2017080887.

23. Lindström, N.O., Sealfon, R., Chen, X., Parvez, R.K., Ransick, A., De Sena Brandine, G., Guo, J., Hill, B., Tran, T., Kim, A.D., et al. (2021). Spatial transcriptional mapping of the human nephrogenic program. Dev. Cell 56, 2381–2398.e6. 10.1016/j.devcel.2021.07.017.

24. Lindström, N.O., Tran, T., Guo, J., Rutledge, E., Parvez, R.K., Thornton, M.E., Grubbs, B., McMahon, J.A., and McMahon, A.P. (2018). Conserved and divergent molecular and anatomic features of human and mouse nephron patterning. J. Am. Soc. Nephrol. 29, 825–840. 10.1681/ASN.2017091036.

25. Lake, B.B., Chen, S., Hoshi, M., Plongthongkum, N., Salamon, D., Knoten, A., Vijayan, A., Venkatesh, R., Kim, E.H., Gao, D., et al. (2019). A single-nucleus RNA-sequencing pipeline to decipher the molecular anatomy and pathophysiology of human kidneys. Nat. Commun. 2019 101 10, 1–15. 10.1038/s41467-019-10861-2.

26. Waterston, R.H., Lindblad-Toh, K., Birney, E., Rogers, J., Abril, J.F., Agarwal, P., Agarwala, R., Ainscough, R., Alexandersson, M., An, P., et al. (2002). Initial sequencing and comparative analysis of the mouse genome. Nature 420, 520–562. 10.1038/nature01262.

27. Lindström, N.O., Guo, J., Kim, A.D., Tran, T., Guo, Q., De Sena Brandine, G., Ransick, A., Parvez, R.K., Thornton, M.E., Baskin, L., et al. (2018). Conserved and Divergent Features of Mesenchymal Progenitor Cell Types within the Cortical Nephrogenic Niche of the Human and Mouse Kidney. J. Am. Soc. Nephrol. 29, 806–824. 10.1681/ASN.2017080890.

28. Potter, E.L. (1965). DEVELOPMENT OF THE HUMAN GLOMERULUS. Arch. Pathol. 80, 241–255.

29. O’Brien, L.L., Guo, Q., Lee, Y.J., Tran, T., Benazet, J.D., Whitney, P.H., Valouev, A., and McMahon, A.P. (2016). Differential regulation of mouse and human nephron progenitors by the six family of transcriptional regulators. Dev. 143, 595–608. 10.1242/dev.127175.

30. Schnell, J., Achieng, M., and Lindström, N.O. (2022). Principles of human and mouse nephron development. Nat. Rev. Nephrol. 10.1038/s41581-022-00598-5.

31. Little, M.H., and McMahon, A.P. (2012). Mammalian kidney development: principles, progress, and projections. Cold Spring Harb. Perspect. Biol. 4. 10.1101/cshperspect.a008300.

32. Combes, A.N. (2015). Towards a quantitative model of kidney morphogenesis. Nephrology 20, 312–314. 10.1111/nep.12407.

33. Hinchliffe, S.A., Sargent, P.H., Howard, C. V, Chan, Y.F., and van Velzen, D. (1991). Human intrauterine renal growth expressed in absolute number of glomeruli assessed by the disector method and Cavalieri principle. Lab. Invest. 64, 777–784.

34. Stuart, T., Butler, A., Hoffman, P., Hafemeister, C., Papalexi, E., Mauck, W.M. 3rd, Hao, Y., Stoeckius, M., Smibert, P., and Satija, R. (2019). Comprehensive Integration of Single-Cell Data. Cell 177, 1888–1902.e21. 10.1016/j.cell.2019.05.031.

35. Buenrostro, J.D., Giresi, P.G., Zaba, L.C., Chang, H.Y., and Greenleaf, W.J. (2013). Transposition of native chromatin for fast and sensitive epigenomic profiling of open chromatin, DNA-binding proteins and nucleosome position. Nat. Methods 10, 1213–1218. 10.1038/nmeth.2688.

36. Stuart, T., Srivastava, A., Madad, S., Lareau, C.A., and Satija, R. (2021). Single-cell chromatin state analysis with Signac. Nat. Methods 18, 1333–1341. 10.1038/s41592-021-01282-5.

37. Schep, A.N., Wu, B., Buenrostro, J.D., and Greenleaf, W.J. (2017). chromVAR: inferring transcription-factor-associated accessibility from single-cell epigenomic data. Nat. Methods 14, 975–978. 10.1038/nmeth.4401.

38. Fornes, O., Castro-Mondragon, J.A., Khan, A., van der Lee, R., Zhang, X., Richmond, P.A., Modi, B.P., Correard, S., Gheorghe, M., Baranašić, D., et al. (2020). JASPAR 2020: update of the open-access database of transcription factor binding profiles. Nucleic Acids Res. 48, D87–D92. 10.1093/nar/gkz1001.

39. Kuhlbrodt, K., Herbarth, B., Sock, E., Hermans-Borgmeyer, I., and Wegner, M. (1998). Sox10, a novel transcriptional modulator in glial cells. J. Neurosci. Off. J. Soc. Neurosci. 18, 237–250. 10.1523/JNEUROSCI.18-01-00237.1998.

40. Nitzan, E., Pfaltzgraff, E.R., Labosky, P.A., and Kalcheim, C. (2013). Neural crest and Schwann cell progenitor-derived melanocytes are two spatially segregated populations similarly regulated by Foxd3. Proc. Natl. Acad. Sci. U. S. A. 110, 12709–12714. 10.1073/pnas.1306287110.

41. Liu, Z., Jin, Y.-Q., Chen, L., Wang, Y., Yang, X., Cheng, J., Wu, W., Qi, Z., and Shen, Z. (2015). Specific marker expression and cell state of Schwann cells during culture in vitro. PLoS One 10, e0123278. 10.1371/journal.pone.0123278.

42. Quesnel-Vallières, M., Irimia, M., Cordes, S.P., and Blencowe, B.J. (2015). Essential roles for the splicing regulator nSR100/SRRM4 during nervous system development. Genes Dev. 29, 746–759. 10.1101/gad.256115.114.

43. Ohnishi, T., Shirane, M., and Nakayama, K.I. (2017). SRRM4-dependent neuron-specific alternative splicing of protrudin transcripts regulates neurite outgrowth. Sci. Rep. 7, 1–13. 10.1038/srep41130.

44. Lo, L., Morin, X., Brunet, J.F., and Anderson, D.J. (1999). Specification of neurotransmitter identity by Phox2 proteins in neural crest stem cells. Neuron 22, 693– 705. 10.1016/s0896-6273(00)80729-1.

45. Pattyn, A., Morin, X., Cremer, H., Goridis, C., and Brunet, J.F. (1999). The homeobox gene Phox2b is essential for the development of autonomic neural crest derivatives. Nature 399, 366–370. 10.1038/20700.

46. Morikawa, Y., D’Autréaux, F., Gershon, M.D., and Cserjesi, P. (2007). Hand2 determines the noradrenergic phenotype in the mouse sympathetic nervous system. Dev. Biol. 307, 114–126. 10.1016/j.ydbio.2007.04.027.

47. Lucas, M.E., Müller, F., Rüdiger, R., Henion, P.D., and Rohrer, H. (2006). The bHLH transcription factor hand2 is essential for noradrenergic differentiation of sympathetic neurons. Development 133, 4015–4024. 10.1242/dev.02574.

48. Bergen, V., Lange, M., Peidli, S., Wolf, F.A., and Theis, F.J. (2020). Generalizing RNA velocity to transient cell states through dynamical modeling. Nat. Biotechnol. 38, 1408– 1414. 10.1038/s41587-020-0591-3.

49. Lawlor, K.T., Zappia, L., Lefevre, J., Park, J.S., Hamilton, N.A., Oshlack, A., Little, M.H., and Combes, A.N. (2019). Nephron progenitor commitment is a stochastic process influenced by cell migration. Elife 8, 1–24. 10.7554/eLife.41156.

50. Matsui, I., Matsumoto, A., Inoue, K., Katsuma, Y., Yasuda, S., Shimada, K., Sakaguchi, Y., Mizui, M., Kaimori, J., Takabatake, Y., et al. (2021). Single cell RNA sequencing uncovers cellular developmental sequences and novel potential intercellular communications in embryonic kidney. Sci. Rep. 11, 73. 10.1038/s41598-020-80154-y.

51. Taguchi, A., Kaku, Y., Ohmori, T., Sharmin, S., Ogawa, M., Sasaki, H., and Nishinakamura, R. (2014). Redefining the in vivo origin of metanephric nephron progenitors enables generation of complex kidney structures from pluripotent stem cells. Cell Stem Cell 14, 53–67. 10.1016/j.stem.2013.11.010.

52. Kuppe, C., Leuchtle, K., Wagner, A., Kabgani, N., Saritas, T., Puelles, V.G., Smeets, B., Hakroush, S., van der Vlag, J., Boor, P., et al. (2019). Novel parietal epithelial cell subpopulations contribute to focal segmental glomerulosclerosis and glomerular tip lesions. Kidney Int. 96, 80–93. 10.1016/j.kint.2019.01.037.

53. Trapnell, C., Cacchiarelli, D., Grimsby, J., Pokharel, P., Li, S., Morse, M., Lennon, N.J., Livak, K.J., Mikkelsen, T.S., and Rinn, J.L. (2014). The dynamics and regulators of cell fate decisions are revealed by pseudotemporal ordering of single cells. Nat. Biotechnol. 32, 381–386. 10.1038/nbt.2859.

54. Barker, N., Rookmaaker, M.B., Kujala, P., Ng, A., Leushacke, M., Snippert, H., van de Wetering, M., Tan, S., Van Es, J.H., Huch, M., et al. (2012). Lgr5(+ve) stem/progenitor cells contribute to nephron formation during kidney development. Cell Rep. 2, 540–552. 10.1016/j.celrep.2012.08.018.

55. Tran, T., Lindström, N.O., Ransick, A., De Sena Brandine, G., Guo, Q., Kim, A.D., Der, B., Peti-Peterdi, J., Smith, A.D., Thornton, M., et al. (2019). In Vivo Developmental Trajectories of Human Podocyte Inform In Vitro Differentiation of Pluripotent Stem Cell-Derived Podocytes. Dev. Cell 50, 102–116.e6. 10.1016/j.devcel.2019.06.001.

56. Grieshammer, U., Cebrián, C., Ilagan, R., Meyers, E., Herzlinger, D., and Martin, G.R. (2005). FGF8 is required for cell survival at distinct stages of nephrogenesis and for regulation of gene expression in nascent nephrons. Development 132, 3847–3857. 10.1242/dev.01944.

57. Butler, A., Hoffman, P., Smibert, P., Papalexi, E., and Satija, R. (2018). Integrating single-cell transcriptomic data across different conditions, technologies, and species. Nat. Biotechnol. 36, 411–420. 10.1038/nbt.4096.

58. Xue, Z., Huang, K., Cai, C., Cai, L., Jiang, C.Y., Feng, Y., Liu, Z., Zeng, Q., Cheng, L., Sun, Y.E., et al. (2013). Genetic programs in human and mouse early embryos revealed by single-cell RNA sequencing. Nature 500, 593–597. 10.1038/nature12364.

59. Kuleshov, M. V, Jones, M.R., Rouillard, A.D., Fernandez, N.F., Duan, Q., Wang, Z., Koplev, S., Jenkins, S.L., Jagodnik, K.M., Lachmann, A., et al. (2016). Enrichr: a comprehensive gene set enrichment analysis web server 2016 update. Nucleic Acids Res. 44, W90–7. 10.1093/nar/gkw377.

60. Chen, E.Y., Tan, C.M., Kou, Y., Duan, Q., Wang, Z., Meirelles, G.V., Clark, N.R., and Ma’ayan, A. (2013). Enrichr: interactive and collaborative HTML5 gene list enrichment analysis tool. BMC Bioinformatics 14, 128. 10.1186/1471-2105-14-128.

61. Binder, J.X., Pletscher-Frankild, S., Tsafou, K., Stolte, C., O’Donoghue, S.I., Schneider, R., and Jensen, L.J. (2014). COMPARTMENTS: unification and visualization of protein subcellular localization evidence. Database (Oxford). 2014, bau012. 10.1093/database/bau012.

62. Kuhn, R.M., Haussler, D., and Kent, W.J. (2013). The UCSC genome browser and associated tools. Brief. Bioinform. 14, 144–161. 10.1093/bib/bbs038.

63. Moore, J.E., Purcaro, M.J., Pratt, H.E., Epstein, C.B., Shoresh, N., Adrian, J., Kawli, T., Davis, C.A., Dobin, A., Kaul, R., et al. (2020). Expanded encyclopaedias of DNA elements in the human and mouse genomes. Nature 583, 699–710. 10.1038/s41586-020-2493-4.

64. McLean, C.Y., Bristor, D., Hiller, M., Clarke, S.L., Schaar, B.T., Lowe, C.B., Wenger, A.M., and Bejerano, G. (2010). GREAT improves functional interpretation of cis-regulatory regions. Nat. Biotechnol. 28, 495–501. 10.1038/nbt.1630.

65. Cotney, J., Leng, J., Yin, J., Reilly, S.K., DeMare, L.E., Emera, D., Ayoub, A.E., Rakic, P., and Noonan, J.P. (2013). The Evolution of Lineage-Specific Regulatory Activities in the Human Embryonic Limb. Cell 154, 185–196. 10.1016/j.cell.2013.05.056.

66. Shibata, Y., Sheffield, N.C., Fedrigo, O., Babbitt, C.C., Wortham, M., Tewari, A.K., London, D., Song, L., Lee, B.-K., Iyer, V.R., et al. (2012). Extensive Evolutionary Changes in Regulatory Element Activity during Human Origins Are Associated with Altered Gene Expression and Positive Selection. PLOS Genet. 8, e1002789.

67. Schraders, M., Oostrik, J., Huygen, P.L.M., Strom, T.M., van Wijk, E., Kunst, H.P.M., Hoefsloot, L.H., Cremers, C.W.R.J., Admiraal, R.J.C., and Kremer, H. (2010). Mutations in PTPRQ are a cause of autosomal-recessive nonsyndromic hearing impairment DFNB84 and associated with vestibular dysfunction. Am. J. Hum. Genet. 86, 604–610. 10.1016/j.ajhg.2010.02.015.

68. Seifert, R.A., Coats, S.A., Oganesian, A., Wright, M.B., Dishmon, M., Booth, C.J., Johnson, R.J., Alpers, C.E., and Bowen-Pope, D.F. (2003). PTPRQ is a novel phosphatidylinositol phosphatase that can be expressed as a cytoplasmic protein or as a subcellularly localized receptor-like protein. Exp. Cell Res. 287, 374–386. 10.1016/s0014-4827(03)00121-6.

69. Kvon, E.Z., Waymack, R., Gad, M., and Wunderlich, Z. (2021). Enhancer redundancy in development and disease. Nat. Rev. Genet. 22, 324–336. 10.1038/s41576-020-00311-x.

70. Osterwalder, M., Barozzi, I., Tissiéres, V., Fukuda-Yuzawa, Y., Mannion, B.J., Afzal, S.Y., Lee, E.A., Zhu, Y., Plajzer-Frick, I., Pickle, C.S., et al. (2018). Enhancer redundancy provides phenotypic robustness in mammalian development. Nature 554, 239–243. 10.1038/nature25461.

71. Ma, S., Zhang, B., LaFave, L., Chiang, Z., Hu, Y., Ding, J., Brack, A., Kartha, V.K., Law, T., Lareau, C., et al. (2020). Chromatin potential identified by shared single cell profiling of RNA and chromatin. bioRxiv, 2020.06.17.156943. 10.1101/2020.06.17.156943.

72. Chen, L., Bao, Y., Jiang, S., and Zhong, X.B. (2020). The roles of long noncoding rnas hnf1α-as1 and hnf4α-as1 in drug metabolism and human diseases. Non-coding RNA 6, 1–21. 10.3390/ncrna6020024.

73. Daugherty, A.C., Yeo, R.W., Buenrostro, J.D., Greenleaf, W.J., Kundaje, A., and Brunet, A. (2017). Chromatin accessibility dynamics reveal novel functional enhancers in C. elegans. Genome Res. 27, 2096–2107. 10.1101/gr.226233.117.

74. Kuhn, M. (2008). Building Predictive Models in R Using the caret Package. J. Stat. Softw. 28, 1–26. 10.18637/jss.v028.i05.

75. Buniello, A., MacArthur, J.A.L., Cerezo, M., Harris, L.W., Hayhurst, J., Malangone, C., McMahon, A., Morales, J., Mountjoy, E., Sollis, E., et al. (2019). The NHGRI-EBI GWAS Catalog of published genome-wide association studies, targeted arrays and summary statistics 2019. Nucleic Acids Res. 47, D1005–D1012. 10.1093/nar/gky1120.

76. Xie, J., Liu, L., Mladkova, N., Li, Y., Ren, H., Wang, W., Cui, Z., Lin, L., Hu, X., Yu, X., et al. (2020). The genetic architecture of membranous nephropathy and its potential to improve non-invasive diagnosis. Nat. Commun. 11, 1600. 10.1038/s41467-020-15383-w.

77. Waterston, R.H., Lindblad-Toh, K., Birney, E., Rogers, J., Abril, J.F., Agarwal, P., Agarwala, R., Ainscough, R., Alexandersson, M., An, P., et al. (2002). Initial sequencing and comparative analysis of the mouse genome. Nature 420, 520–562. 10.1038/nature01262.

78. Hay, M., Thomas, D.W., Craighead, J.L., Economides, C., and Rosenthal, J. (2014). Clinical development success rates for investigational drugs. Nat. Biotechnol. 32, 40–51. 10.1038/nbt.2786.

79. Breschi, A., Gingeras, T.R., and Guigó, R. (2017). Comparative transcriptomics in human and mouse. Nat. Rev. Genet. 18, 425–440. 10.1038/nrg.2017.19.

80. Combes, A.N., Phipson, B., Lawlor, K.T., Dorison, A., Patrick, R., Zappia, L., Harvey, R.P., Oshlack, A., and Little, M.H. (2019). Single cell analysis of the developing mouse kidney provides deeper insight into marker gene expression and ligand-receptor crosstalk. Development 146. 10.1242/dev.178673.

81. Muto, Y., Wilson, P.C., Ledru, N., Wu, H., Dimke, H., Waikar, S.S., and Humphreys, B.D. (2021). Single cell transcriptional and chromatin accessibility profiling redefine cellular heterogeneity in the adult human kidney. Nat. Commun. 2021 121 12, 1–17. 10.1038/s41467-021-22368-w.

82. Chen, L., Lee, J.W., Chou, C.L., Nair, A. V., Battistone, M.A., Păunescu, T.G., Merkulova, M., Breton, S., Verlander, J.W., Wall, S.M., et al. (2017). Transcriptomes of major renal collecting duct cell types in mouse identified by single-cell RNA-seq. Proc. Natl. Acad. Sci. U. S. A. 114, E9989–E9998. 10.1073/pnas.1710964114.

83. Magella, B., Adam, M., Potter, A.S., Venkatasubramanian, M., Chetal, K., Hay, S.B., Salomonis, N., and Potter, S.S. (2018). Cross-platform single cell analysis of kidney development shows stromal cells express Gdnf. Dev. Biol. 434, 36–47. 10.1016/j.ydbio.2017.11.006.

84. Zimmerman, K.A., Bentley, M.R., Lever, J.M., Li, Z., Crossman, D.K., Song, C.J., Liu, S., Crowley, M.R., George, J.F., Mrug, M., et al. (2019). Single-cell RNA sequencing identifies candidate renal resident macrophage gene expression signatures across species. J. Am. Soc. Nephrol. 30, 767–781. 10.1681/ASN.2018090931.

85. Wang, P., Chen, Y., Yong, J., Cui, Y., Wang, R., Wen, L., Qiao, J., and Tang, F. (2018). Dissecting the Global Dynamic Molecular Profiles of Human Fetal Kidney Development by Single-Cell RNA Sequencing. Cell Rep. 24, 3554–3567.e3. 10.1016/j.celrep.2018.08.056.

86. Hochane, M., van den Berg, P.R., Fan, X., Bérenger-Currias, N., Adegeest, E., Bialecka, M., Nieveen, M., Menschaart, M., Chuva de Sousa Lopes, S.M., and Semrau, S. (2019). Single-cell transcriptomics reveals gene expression dynamics of human fetal kidney development 10.1371/journal.pbio.3000152.

87. Menon, R., Otto, E.A., Kokoruda, A., Zhou, J., Zhang, Z., Yoon, E., Chen, Y.C., Troyanskaya, O., Spence, J.R., Kretzler, M., et al. (2018). Single-cell analysis of progenitor cell dynamics and lineage specification in the human fetal kidney. Dev. 145. 10.1242/dev.164038.

88. Rowe, R.G., and Daley, G.Q. (2019). Induced pluripotent stem cells in disease modelling and drug discovery. Nat. Rev. Genet. 20, 377–388. 10.1038/s41576-019-0100-z.

89. Meerabux, J.M.A., Ohba, H., Fukasawa, M., Suto, Y., Aoki-Suzuki, M., Nakashiba, T., Nishimura, S., Itohara, S., and Yoshikawa, T. (2005). Human netrin-G1 isoforms show evidence of differential expression. Genomics 86, 112–116. 10.1016/j.ygeno.2005.04.004.

90. Tran, T., Song, C.J., Nguyen, T., Cheng, S.-Y., McMahon, J.A., Yang, R., Guo, Q., Der, B., Lindström, N.O., Lin, D.C.-H., et al. (2022). A scalable organoid model of human autosomal dominant polycystic kidney disease for disease mechanism and drug discovery. Cell Stem Cell 29, 1083–1101.e7. 10.1016/j.stem.2022.06.005.

91. Hart, T.C., Gorry, M.C., Hart, P.S., Woodard, A.S., Shihabi, Z., Sandhu, J., Shirts, B., Xu, L., Zhu, H., Barmada, M.M., et al. (2002). Mutations of the UMOD gene are responsible for medullary cystic kidney disease 2 and familial juvenile hyperuricaemic nephropathy. J. Med. Genet. 39, 882–892. 10.1136/jmg.39.12.882.

92. Wuttke, M., Li, Y., Li, M., Sieber, K.B., Feitosa, M.F., Gorski, M., Tin, A., Wang, L., Chu, A.Y., Hoppmann, A., et al. (2019). A catalog of genetic loci associated with kidney function from analyses of a million individuals. Nat. Genet. 51, 957–972. 10.1038/s41588-019-0407-x.

93. Köttgen, A., Pattaro, C., Böger, C.A., Fuchsberger, C., Olden, M., Glazer, N.L., Parsa, A., Gao, X., Yang, Q., Smith, A. V, et al. (2010). New loci associated with kidney function and chronic kidney disease. Nat. Genet. 42, 376–384. 10.1038/ng.568.

94. van der Ven, A.T., Connaughton, D.M., Ityel, H., Mann, N., Nakayama, M., Chen, J., Vivante, A., Hwang, D.-Y., Schulz, J., Braun, D.A., et al. (2018). Whole-Exome Sequencing Identifies Causative Mutations in Families with Congenital Anomalies of the Kidney and Urinary Tract. J. Am. Soc. Nephrol. 29, 2348–2361. 10.1681/ASN.2017121265.

95. Subramanian, A., Sidhom, E.-H., Emani, M., Vernon, K., Sahakian, N., Zhou, Y., Kost- Alimova, M., Slyper, M., Waldman, J., Dionne, D., et al. (2019). Single cell census of human kidney organoids shows reproducibility and diminished off-target cells after transplantation. Nat. Commun. 10, 5462. 10.1038/s41467-019-13382-0.

96. Lan, Y., Wang, Q., Ovitt, C.E., and Jiang, R. (2007). A unique mouse strain expressing Cre recombinase for tissue-specific analysis of gene function in palate and kidney development. Genesis 45, 618–624. 10.1002/dvg.20334.

97. D’Agati, V.D., and Shankland, S.J. (2019). Recognizing diversity in parietal epithelial cells. Kidney Int. 96, 16–19. 10.1016/j.kint.2019.02.036.

98. Nakai, S., Sugitani, Y., Sato, H., Ito, S., Miura, Y., Ogawa, M., Nishi, M., Jishage, K.I., Minowa, O., and Noda, T. (2003). Crucial roles fo Brn1 in distal tubule formation and function in mouse kidney. Development 130, 4751–4759. 10.1242/dev.00666.

99. Marable, S.S., Chung, E., and Park, J.S. (2020). Hnf4a is required for the development of cdh6-expressing progenitors into proximal tubules in the mouse kidney. J. Am. Soc. Nephrol. 31, 2543–2558. 10.1681/ASN.2020020184.

100. Wu, H., Kirita, Y., Donnelly, E.L., and Humphreys, B.D. (2019). Advantages of Single-Nucleus over Single-Cell RNA Sequencing of Adult Kidney: Rare Cell Types and Novel Cell States Revealed in Fibrosis. J. Am. Soc. Nephrol. 30, 23–32. 10.1681/ASN.2018090912.

101. Liu, Y., Beyer, A., and Aebersold, R. (2016). On the Dependency of Cellular Protein Levels on mRNA Abundance. Cell 165, 535–550. https://doi.org/10.1016/j.cell.2016.03.014.

102. Hafemeister, C., and Satija, R. (2019). Normalization and variance stabilization of single-cell RNA-seq data using regularized negative binomial regression. Genome Biol. 20, 1–15. 10.1186/s13059-019-1874-1.

103. Zhou, Q., Liu, M., Xia, X., Gong, T., Feng, J., Liu, W., Liu, Y., Zhen, B., Wang, Y., Ding, C., et al. (2017). A mouse tissue transcription factor atlas. Nat. Commun. 8, 15089. 10.1038/ncomms15089.

104. Ng, A.H.M., Khoshakhlagh, P., Rojo Arias, J.E., Pasquini, G., Wang, K., Swiersy, A., Shipman, S.L., Appleton, E., Kiaee, K., Kohman, R.E., et al. (2021). A comprehensive library of human transcription factors for cell fate engineering. Nat. Biotechnol. 39, 510–519. 10.1038/s41587-020-0742-6.

105. Muto, Y., Wilson, P.C., Ledru, N., Wu, H., Dimke, H., Waikar, S.S., and Humphreys, B.D. (2021). Single cell transcriptional and chromatin accessibility profiling redefine cellular heterogeneity in the adult human kidney. Nat. Commun. 12, 2190. 10.1038/s41467-021-22368-w.

106. England, A.R., Chaney, C.P., Das, A., Patel, M., Malewska, A., Armendariz, D., Hon, G.C., Strand, D.W., Drake, K.A., and Carroll, T.J. (2020). Identification and characterization of cellular heterogeneity within the developing renal interstitium. Development 147. 10.1242/dev.190108.

107. Howe, K.L., Achuthan, P., Allen, J., Allen, J., Alvarez-Jarreta, J., Amode, M.R., Armean, I.M., Azov, A.G., Bennett, R., Bhai, J., et al. (2021). Ensembl 2021. Nucleic Acids Res. 49, D884–D891. 10.1093/nar/gkaa942.

108. La Manno, G., Soldatov, R., Zeisel, A., Braun, E., Hochgerner, H., Petukhov, V., Lidschreiber, K., Kastriti, M.E., Lönnerberg, P., Furlan, A., et al. (2018). RNA velocity of single cells. Nature 560, 494–498. 10.1038/s41586-018-0414-6.

109. Virshup, I., Rybakov, S., Theis, F.J., Angerer, P., and Wolf, F.A. (2021). anndata: Annotated data. bioRxiv, 2021.12.16.473007. 10.1101/2021.12.16.473007.

110. Qiu, X., Mao, Q., Tang, Y., Wang, L., Chawla, R., Pliner, H.A., and Trapnell, C. (2017). Reversed graph embedding resolves complex single-cell trajectories. Nat. Methods 14, 979–982. 10.1038/nmeth.4402.

111. Korsunsky, I., Millard, N., Fan, J., Slowikowski, K., Zhang, F., Wei, K., Baglaenko, Y., Brenner, M., Loh, P., and Raychaudhuri, S. (2019). Fast, sensitive and accurate integration of single-cell data with Harmony. Nat. Methods 16, 1289–1296. 10.1038/s41592-019-0619-0.

112. Zhang, Y., Liu, T., Meyer, C.A., Eeckhoute, J., Johnson, D.S., Bernstein, B.E., Nusbaum, C., Myers, R.M., Brown, M., Li, W., et al. (2008). Model-based analysis of ChIP-Seq (MACS). Genome Biol. 9, R137. 10.1186/gb-2008-9-9-r137.

113. Danecek, P., Bonfield, J.K., Liddle, J., Marshall, J., Ohan, V., Pollard, M.O., Whitwham, A., Keane, T., McCarthy, S.A., Davies, R.M., et al. (2021). Twelve years of SAMtools and BCFtools. Gigascience 10. 10.1093/gigascience/giab008.

114. Yu, G., Wang, L.-G., and He, Q.-Y. (2015). ChIPseeker: an R/Bioconductor package for ChIP peak annotation, comparison and visualization. Bioinformatics 31, 2382–2383. 10.1093/bioinformatics/btv145.

115. Ramírez, F., Dündar, F., Diehl, S., Grüning, B.A., and Manke, T. (2014). deepTools: a flexible platform for exploring deep-sequencing data. Nucleic Acids Res. 42, W187–91. 10.1093/nar/gku365.

116. Hart, R.P. (2019). Strategies for Integrating Single-Cell RNA Sequencing Results With Multiple Species. bioRxiv, 671115. 10.1101/671115.

117. Lawrence, M., Huber, W., Pagès, H., Aboyoun, P., Carlson, M., Gentleman, R., Morgan, M.T., and Carey, V.J. (2013). Software for computing and annotating genomic ranges. PLoS Comput. Biol. 9, e1003118. 10.1371/journal.pcbi.1003118.

118. Ma, S., Zhang, B., LaFave, L.M., Earl, A.S., Chiang, Z., Hu, Y., Ding, J., Brack, A., Kartha, V.K., Tay, T., et al. (2020). Chromatin Potential Identified by Shared Single-Cell Profiling of RNA and Chromatin. Cell 183, 1103–1116.e20. 10.1016/j.cell.2020.09.056.

119. Pliner, H.A., Packer, J.S., McFaline-Figueroa, J.L., Cusanovich, D.A., Daza, R.M., Aghamirzaie, D., Srivatsan, S., Qiu, X., Jackson, D., Minkina, A., et al. (2018). Cicero Predicts cis-Regulatory DNA Interactions from Single-Cell Chromatin Accessibility Data. Mol. Cell 71, 858–871.e8. 10.1016/j.molcel.2018.06.044.

120. Liaw, A., and Wiener, M. (2002). Classification and Regression by randomForest. R News 2, 18–22.

121. van der Ven, A.T., Vivante, A., and Hildebrandt, F. (2018). Novel Insights into the Pathogenesis of Monogenic Congenital Anomalies of the Kidney and Urinary Tract. J. Am. Soc. Nephrol. 29, 36–50. 10.1681/ASN.2017050561.

