## Supplementary figures and images for "Comparative single-cell analyses identify shared and divergent features of human and mouse kidney development"

### Supplementary figure 1

Supplementary figure S1.

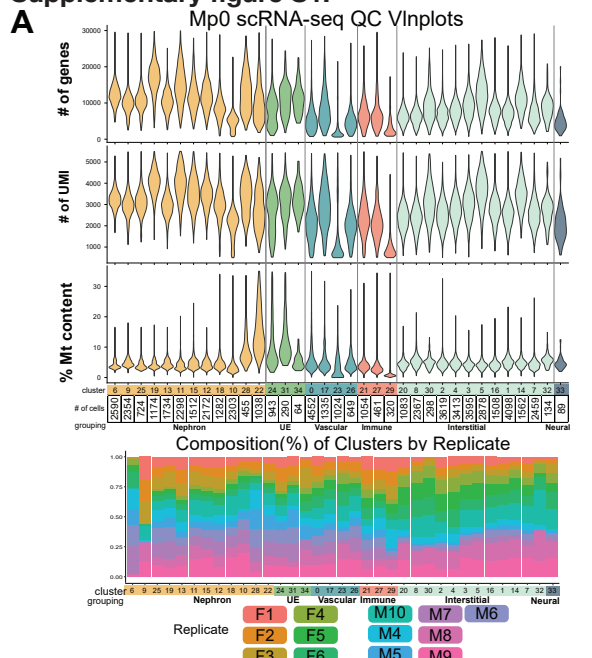

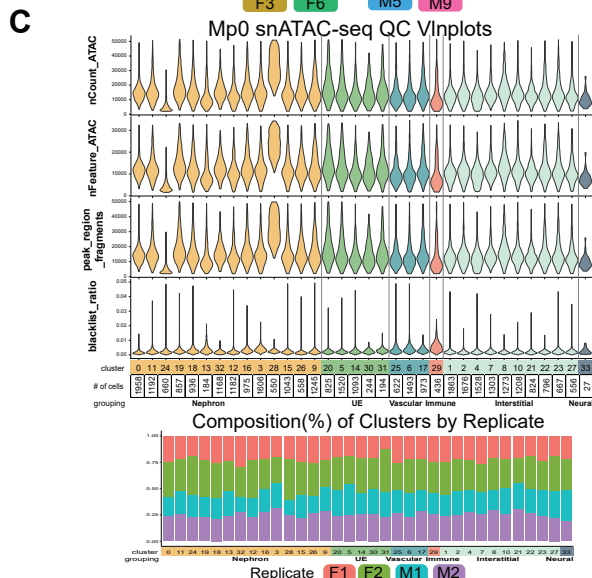

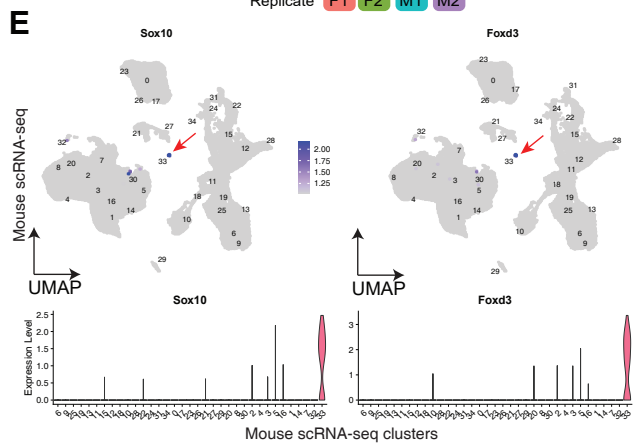

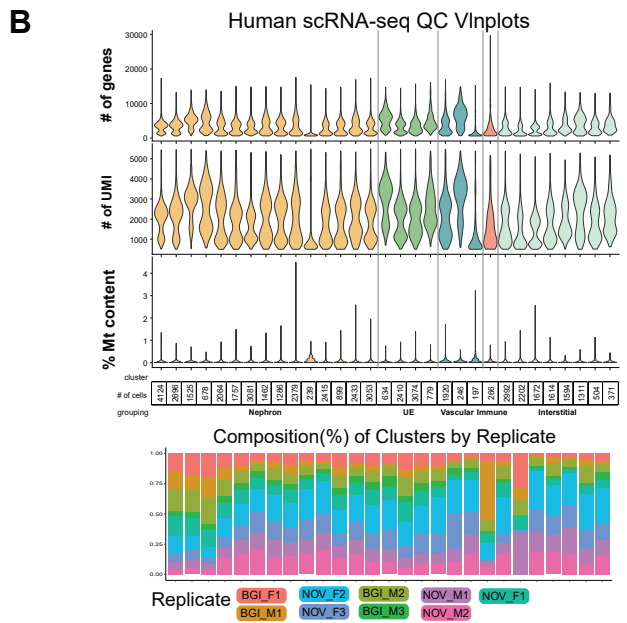

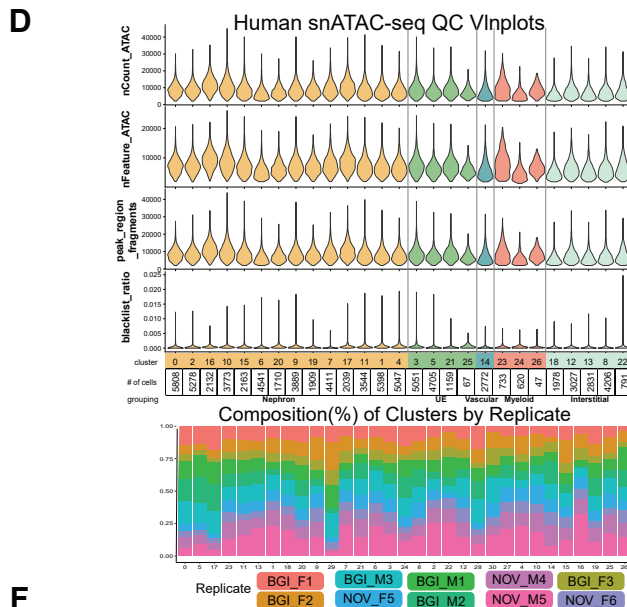

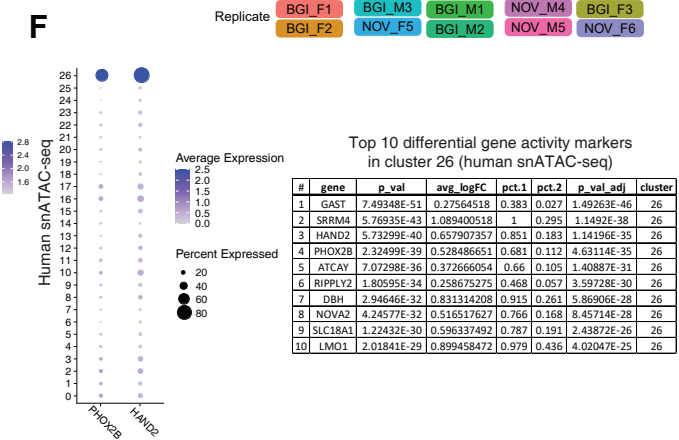

### Supplementary figure 3

#### Supplementary figure S3.

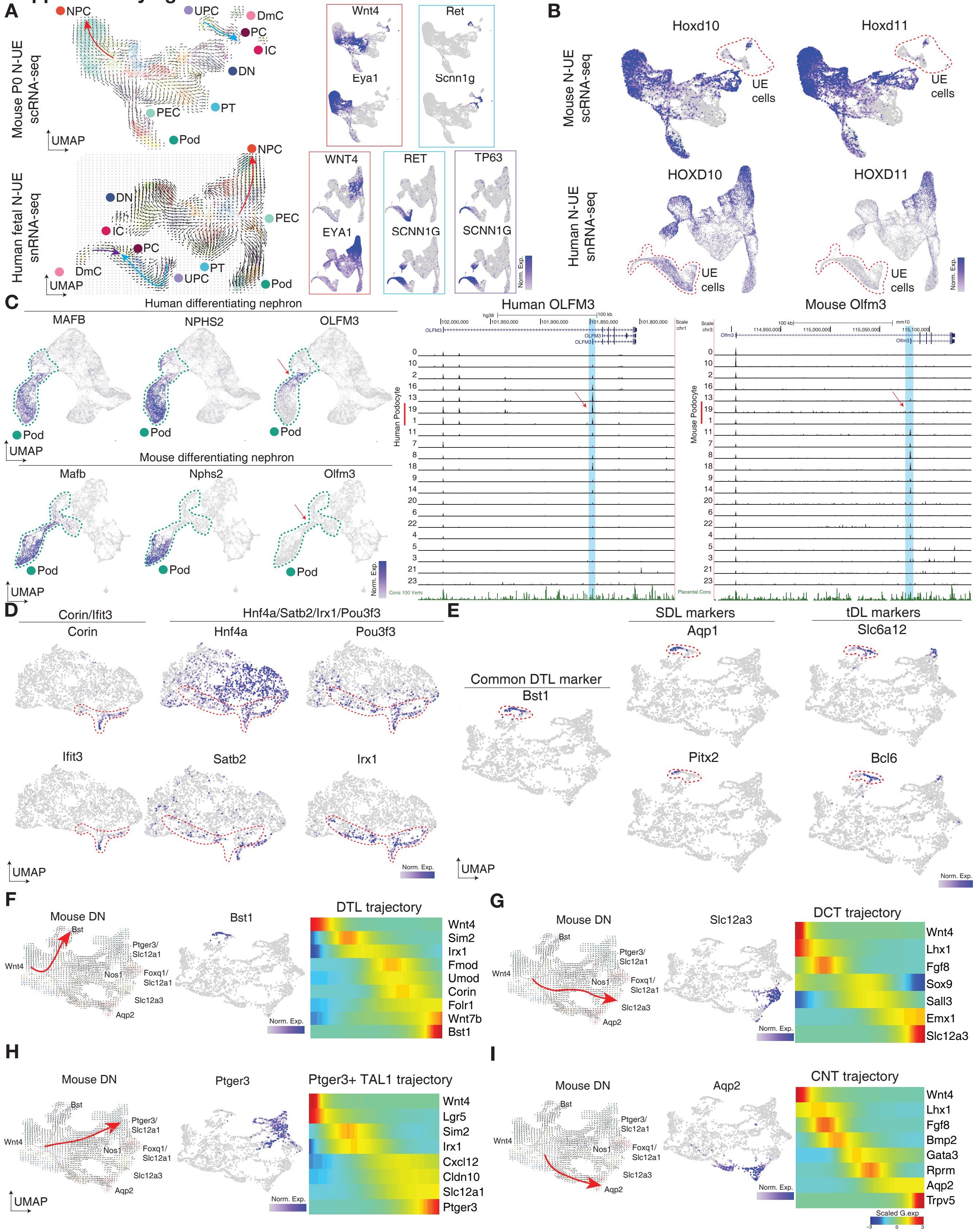

### Supplementary figure 4

#### Supplementary figure S4.

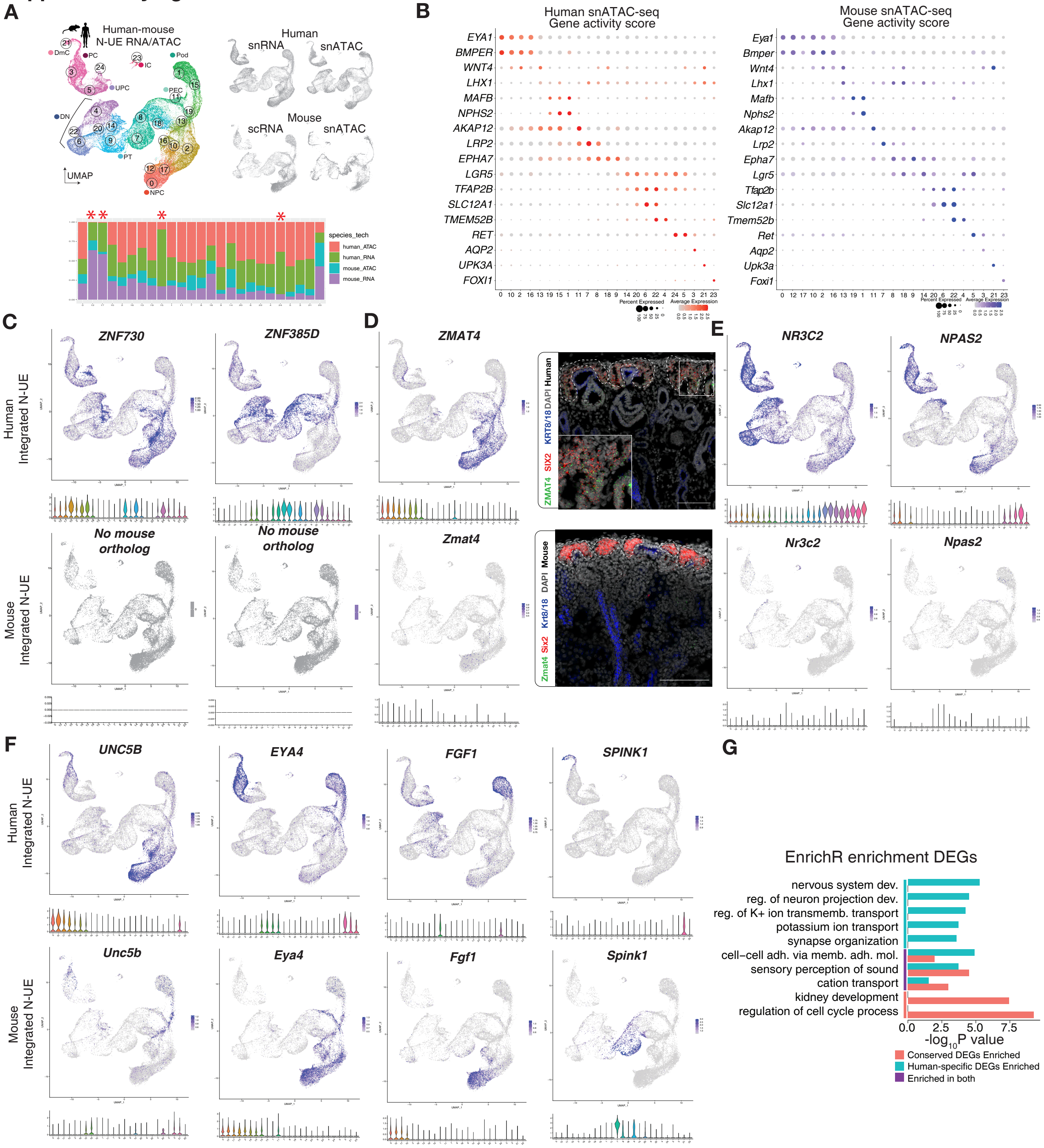

### Supplementary figure 5

#### Supplementary figure S5.

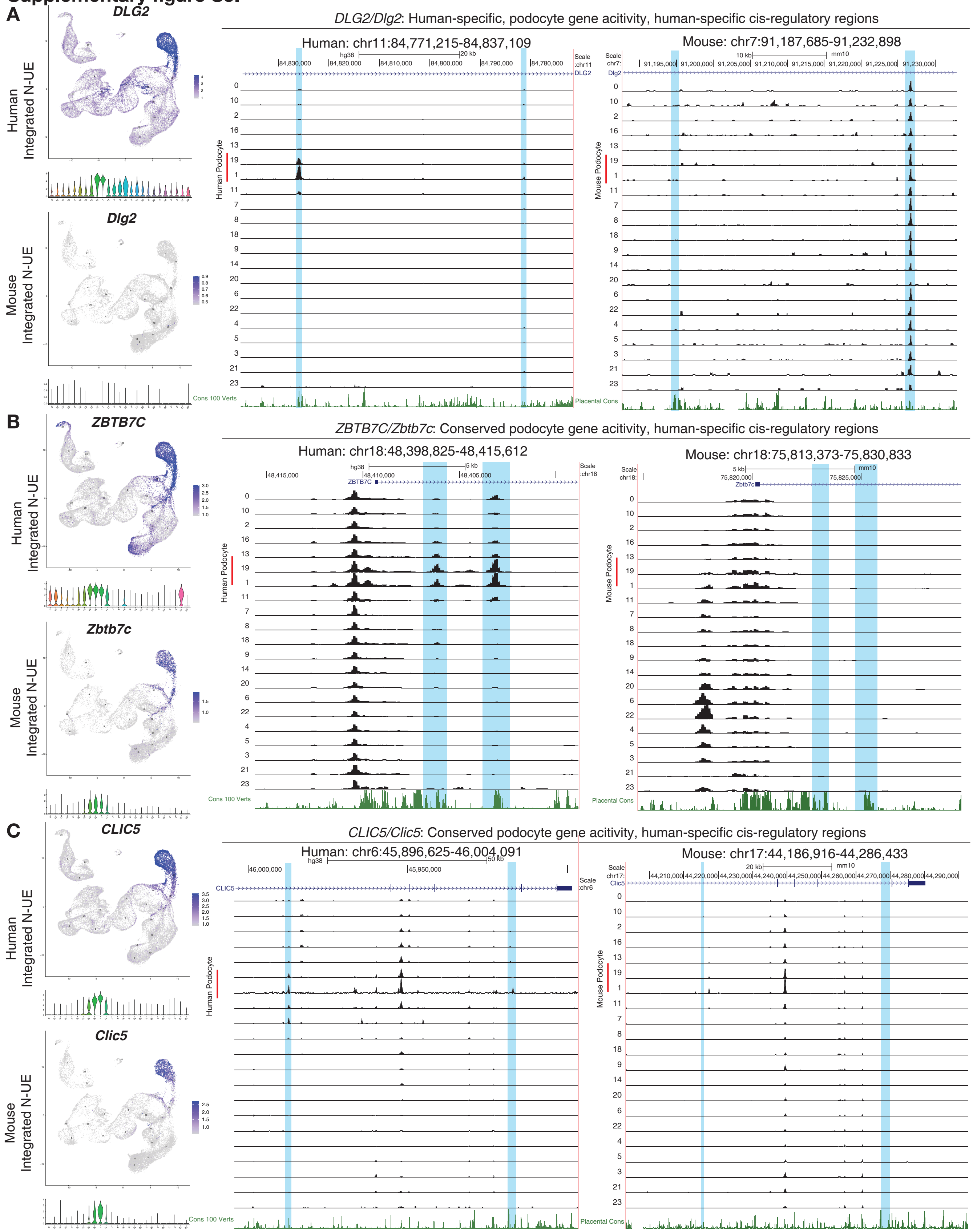

### Supplementary figure 6

**Supplementary figure S6.**

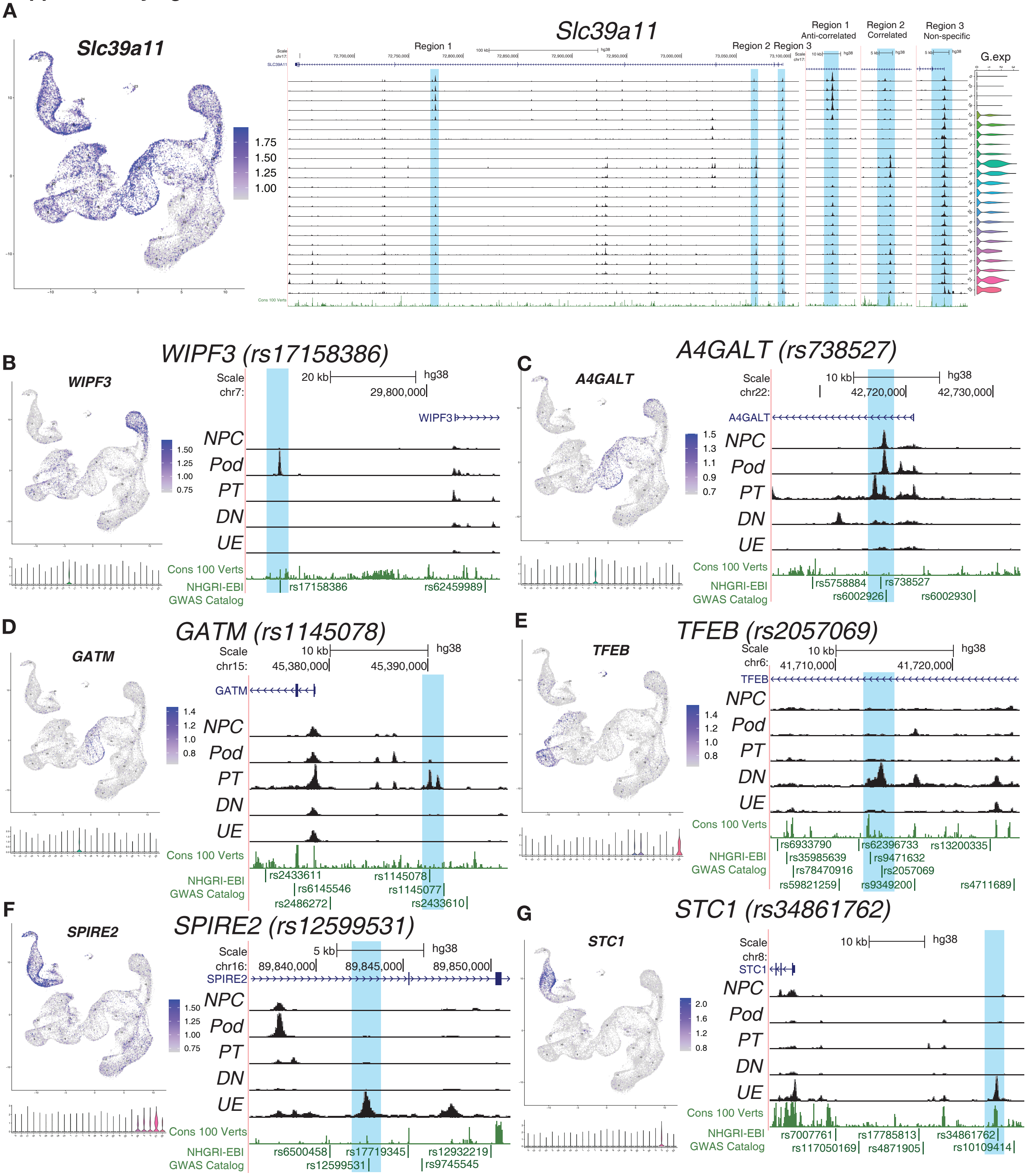
