## Supplementary figure 2 for "Comparative single-cell analyses identify shared and divergent features of human and mouse kidney development"

### Supplementary figure S2.

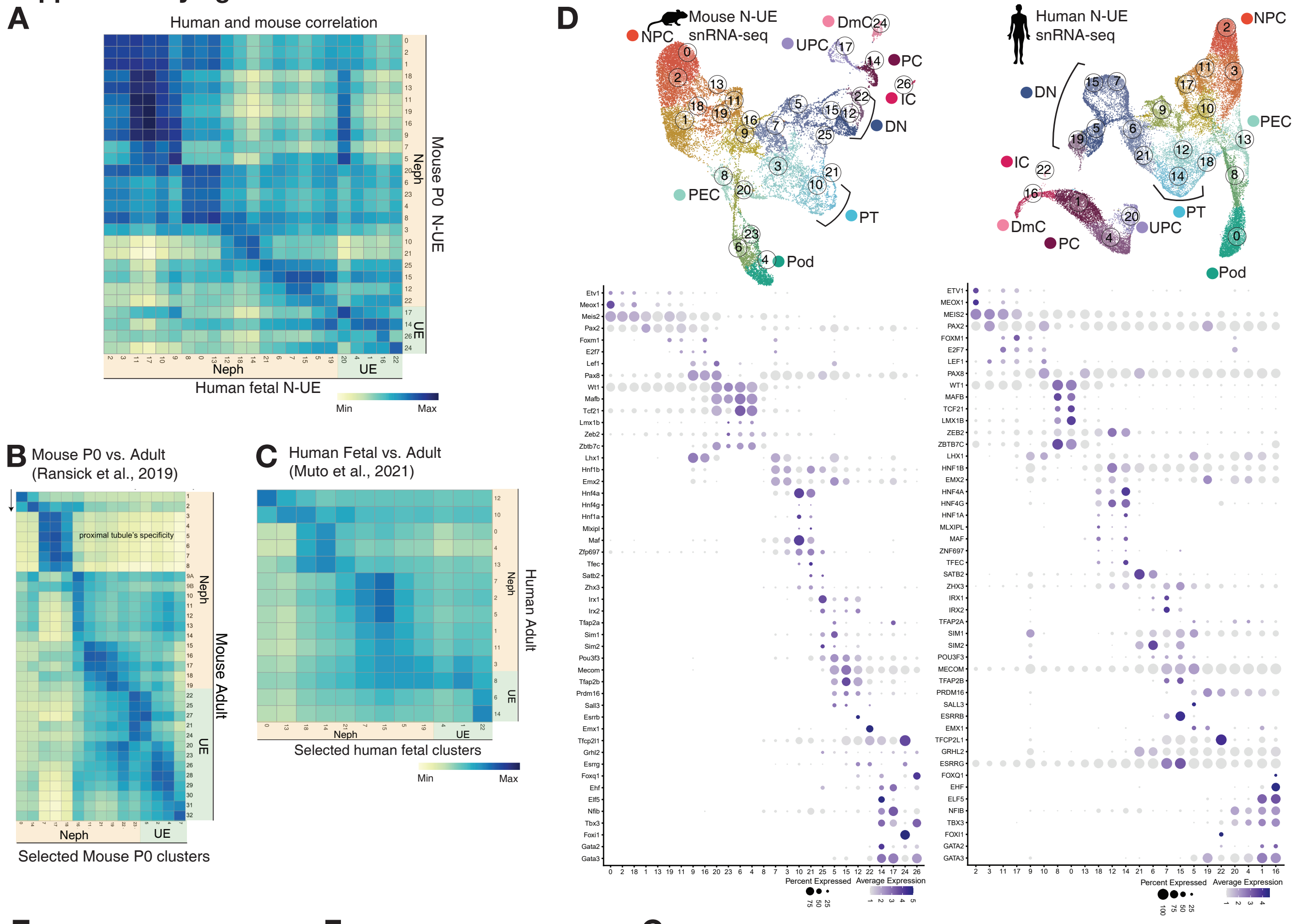


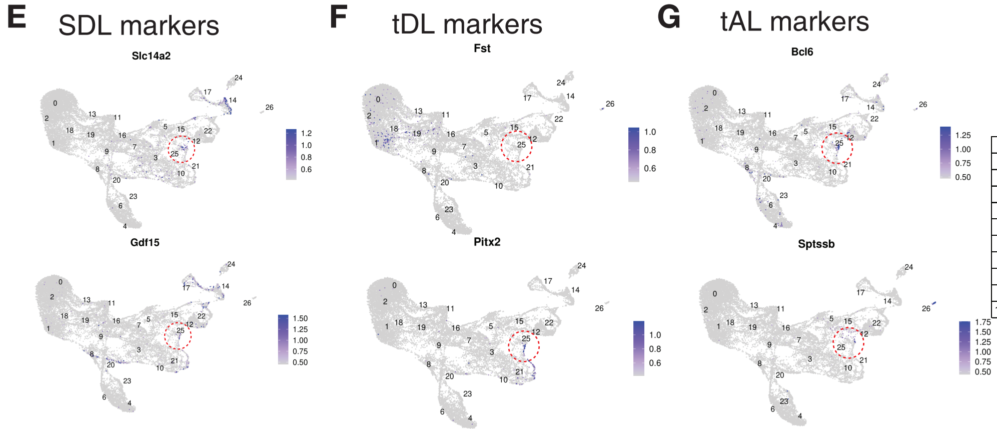


Top 10 markers in cluster 25
(Mouse N-UE scRNA-seq)

| # | gene | avg_logFC | pct.1 | pct.2 | p_val_adj | cluster |
|---|---|---|---|---|---|---|
| 1 | Ifit3 | 1.415 | 0.607 | 0.021 | 0 | 25 |
| 2 | Irx1 | 1.273 | 0.737 | 0.067 | 0 | 25 |
| 3 | Corin | 1.046 | 0.629 | 0.029 | 0 | 25 |
| 4 | Bst1 | 0.903 | 0.272 | 0.008 | 0 | 25 |
| 5 | D030045P18Rik | 0.856 | 0.701 | 0.063 | 0 | 25 |
| 6 | Cfap52 | 0.776 | 0.366 | 0.01 | 0 | 25 |
| 7 | Wnt7b | 0.710 | 0.509 | 0.032 | 0 | 25 |
| 8 | Ifit3b | 0.527 | 0.312 | 0.007 | 0 | 25 |
| 9 | Umod | 1.213 | 0.603 | 0.049 | 1.68E-287 | 25 |
| 10 | Slc6a12 | 0.424 | 0.196 | 0.005 | 2.20E-264 | 25 |
